# Quorum sensing mediates morphology and motility transitions in the model archaeon *Haloferax volcanii*

**DOI:** 10.1101/2025.01.14.633064

**Authors:** Priyanka Chatterjee, Caroline E. Consoli, Heather Schiller, Kiersten K. Winter, Monica E. McCallum, Stefan Schulze, Mechthild Pohlschroder

## Abstract

Quorum sensing (QS) is a mechanism of intercellular communication that enables microbes to alter gene expression and adapt to the environment. This cell-cell signaling is necessary for intra- and interspecies behaviors such as virulence and biofilm formation. While QS has been extensively studied in bacteria, little is known about cell-cell communication in archaea. Here we established an archaeal model system to study QS. We showed that for *Haloferax volcanii*, the transition from motile rods to non-motile disks is dependent on a possibly novel, secreted small molecule present in cell-free conditioned medium (CM). Moreover, we determined that this putative QS molecule fails to induce the morphology transition in mutants lacking the regulatory factors, DdfA and CirA. Using quantitative proteomics of wild-type cells, we detected significant differential abundances of 236 proteins in the presence of CM. Conversely, in the Δ*ddfA* mutant, addition of CM resulted in only 110 proteins of significant differential abundances. These results confirm that DdfA is involved in CM-dependent regulation. CirA, along with other proteins involved in morphology and swimming motility transitions, is among the proteins regulated by DdfA. These discoveries significantly advance our understanding of microbial communication within archaeal species.

## Main Text Introduction

Microorganisms employ many different forms of intercellular communication critical for survival. The most well-studied mechanism of microbial cell-cell communication is quorum sensing (QS), in which microorganisms synthesize and secrete small molecules or peptides called autoinducers. Recognition of the autoinducers at varying extracellular concentrations enable the microbe to regulate gene expression as a result of changes in their population density. Only when a density threshold is reached, or quorum of molecules is recognized, do the single cells express genes to orchestrate collective behaviors such as bioluminescence, biofilm formation, competence, motility and sporulation^1–4^. Studies about bacterial intercellular communication began with competence factor ComX in *Streptococcus pneumoniae*^5,6^ and the acyl-homoserine lactone (AHL) regulating bioluminescence in *Vibrio* species^7–9^. In subsequent years, many other QS molecules were identified and characterized, such as autoinducer-2 (AI-2)^10,11^, *Pseudomonas* quinolone signal (PQS)^12^, autoinducing peptide (AIP) of gram-positive bacteria^13^, and diffusible signal factor (DSF)^14^.

These QS signaling molecules are synthesized and recognized by different mechanisms. One of the best-defined QS pathways is the LuxI/LuxR system. LuxI catalyzes the synthesis of AHL molecules that can diffuse across the membrane. At a critical extracellular concentration of AHL molecules, signifying that a quorum of bacterial kin is present, AHL can bind to and stabilize the LuxR dimer, which then promotes transcription of QS activated genes^15^. As another example of a QS pathway, in Gram-positive bacteria, QS behaviors can be mediated by two-component systems. Autoinducer molecules bind histidine kinase receptors in the membrane leading to phosphorylation of cognate response regulator proteins. The majority of response regulators in Gram-positive QS pathways function through transcriptional regulation of target genes^16^. In this way, characterizing both the molecules and mechanisms driving microbial intercellular communication enables us to eavesdrop on the microbial chatter, deepening our understanding of single-cellular life as well as providing targets for therapeutics or disinfectants that can interrupt the crosstalk^17^.

In recent years, QS mechanisms have been discovered in non-bacterial microorganisms as well. For the fungus *Candida albicans*, farnesol is an important signal for morphology changes^18^, for the methanogenic archaeon *Methanosaeta harundinacea* 6Ac, a carboxylated AHL regulates cell assembly and carbon metabolic flux^19^, and even for the virus, phi3T phage, the SAIRGA peptide mediates the phage lysis versus lysogeny decision in the host^20^. These discoveries emphasize the evolutionary significance of density-dependent signaling and its fundamental role in microbial life.

Archaea are ubiquitous in the environment^21^, play critical roles in geochemical cycles^22^, and have demonstrated applications in biotechnology^23,24^, yet only a handful of studies have explored the potential for intercellular signaling in archaea. In two bioinformatic studies, putative LuxR solos (LuxR homologs without a cognate LuxI homolog) were identified in many archaeal species^25^, and a strong correlation was observed between metabolic regulation and candidate QS genes in ammonia-oxidizing archaea^26^. Interestingly, multiple studies have revealed that crude supernatant extract from different archaeal species can stimulate AHL-dependent QS phenotypes of bacterial bioreporters, including *M. harundinacea* supernatant containing the carboxy-AHL and *Haloterrigena hispanica* supernatant containing a class of diketopiperazine (DKP) compounds as the active molecules^19,27–30^. These findings suggest the potential for inter- domain crosstalk. However, the identity of the active molecules in most of the archaeal strains tested in the various bioreporter assays remain unknown^27,29,30^, and it is also unknown whether the DKP molecules from *Htr. hispanica* act as true QS signals that induce changes in the archaeon’s behavior^28^. Only three examples of true archaeal QS, or population density-dependent phenotypic changes in archaea mediated by secreted molecules, have been described: *M. harundinacea* morphology changes mediated by carboxy-AHL^19^, growth-phase dependent biosynthesis of *Natrialba magadii* Nep protease (unknown signal molecule)^31^, and increased biofilm biomass of *Halorubrum lacusprofundi* upon the addition of culture supernatant (unknown signal molecule)^29,32^. Furthermore, *M. harundinacea filI* gene is the only gene discovered in any archaeon that has been empirically determined to be involved in a QS pathway^19^. The dearth of information about archaeal QS may be a result of the potential challenges associated with culturing archaea, such as the specific metabolic and environmental needs of extremophilic archaea, as well as the general lack of archaeal genome annotations^33^. Because observing QS-dependent phenotypes in the lab requires microbial cultures, there remains a need for the development of an easily culturable and genetically tractable archaeal model system to study archaeal cell-cell signaling.

The halophilic archaeon *Haloferax volcanii* is easy to culture in the lab and has a wide array of established genetic and biochemical assays^34^. This model archaeon also displays distinct morphology differences at varying population densities, indicating the potential for QS-related behaviors. In liquid culture, cells initially present as motile rods at early log growth phase. As the culture propagates, cells begin to transition to pleomorphic disks, presumably increasing the surface area to volume ratio. By the time the cell population levels off at its highest density, cells are exclusively disk-shaped^35–37^. This morphological change can also be observed in swimming motility halos on soft agar plates, with non-motile disks at the dense center of the halo, and motile rods at the less dense, leading edge^38^.

In this report, we demonstrate that *Hfx. volcanii* is an invaluable model for deciphering QS in archaea. Our interdisciplinary studies strongly suggest that a small molecule, one that has not been previously identified as involved in density-dependent signaling, is secreted by *Hfx. volcanii* to mediate the morphology transition. We identified two components, DdfA and CirA, involved in the QS response and employed the deletion strain Δ*ddfA* in quantitative proteomics to identify additional potential QS components. Thus, by developing methods to analyze a previously uncharacterized QS system in the model archaeon *Hfx. volcanii*, we pave the road for future studies into archaeal cell-cell communication.

## Results

### *Hfx. volcanii* rod-to-disk transition is population density-dependent

To determine if *Hfx. volcanii* disk-formation is mediated by QS molecules secreted into the medium, cell-free supernatant from a stationary phase culture (OD_600_ 1.5) was used to mimic high cell density. Applying this conditioned medium (CM) to fresh *Hfx. volcanii* cultures resulted in disk-shaped, instead of rod-shaped, cells in early log phase (OD_600_ 0.05) (**Fig. 1a**). As little as 1% CM resulted in the early log cultures lacking rods entirely (**Fig. 1b, S1**), strongly suggesting that the replacement of fresh media, or nutrient depletion, is not the cause of the shape change. Consistent with the hypothesis that *Hfx. volcanii* shape transition is population density-dependent, addition of CM from cultures grown to higher OD_600_ correlated to higher percentages of disk-shaped cells at early log phase (**Fig. 1c**).

**Figure 1.**
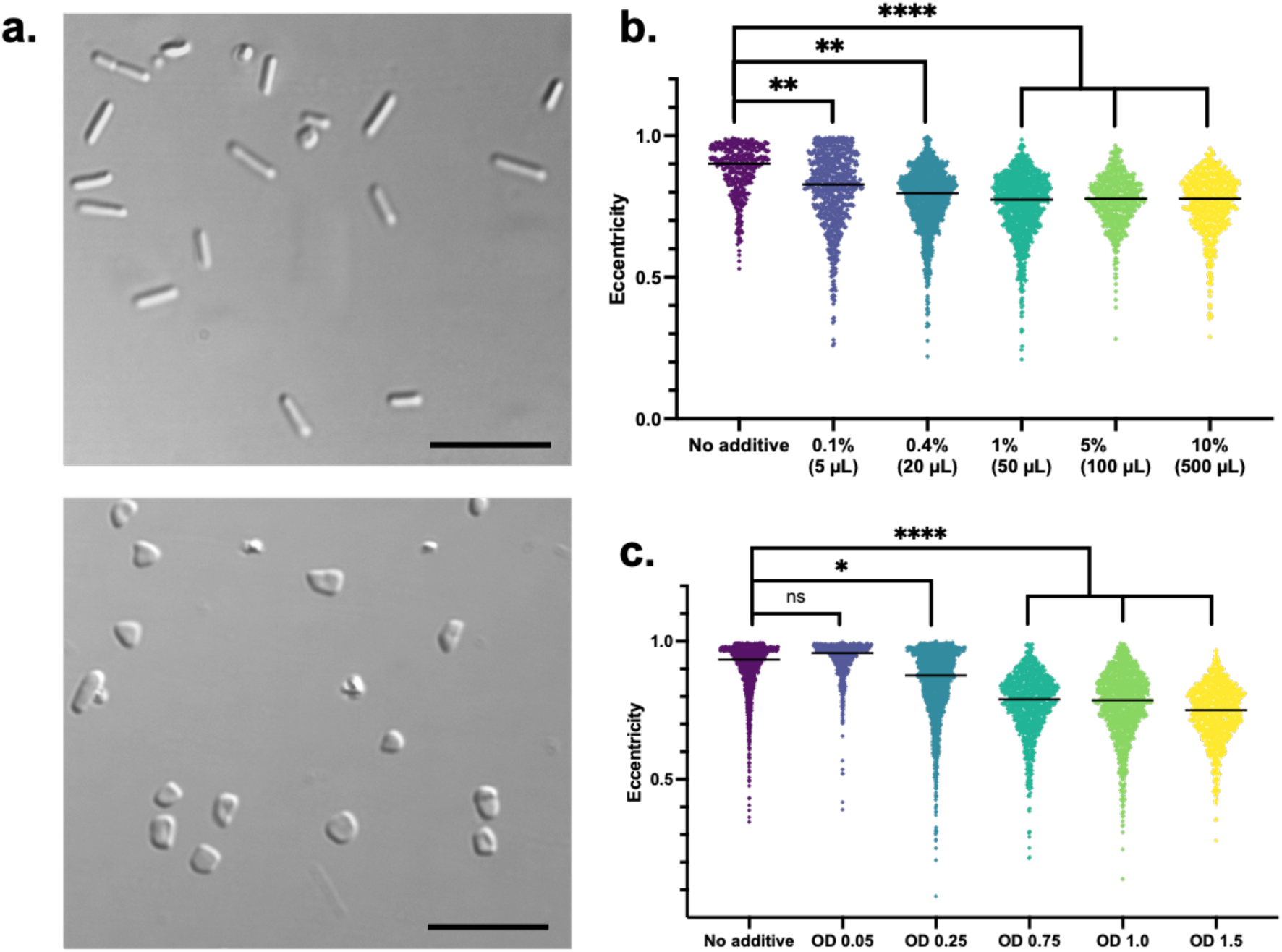
*Hfx. volcanii* cells are disk-shaped in early log in the presence of conditioned medium (CM). Differential interference contrast (DIC) images of *Hfx. volcanii* cells from 5 mL cultures grown to OD600 0.05 with no additive (top) and with 50 µL of CM (1% v/v) added (bottom). Scale bar is 10 µm **(a)**. Cell shape quantification of three biological replicates of each condition to determine lowest effective volume of CM from an *Hfx. volcanii* culture grown to OD600 1.5 **(b)** or of the effect of 1% CM from cultures grown to differing optical densities **(c)** by using the eccentricity parameter of CellProfiler^55,56^, which outputs a value between 0 (circle) and 1 (line segment). Eccentricity values that range between 0.5 and 0.9 correspond to disks while values that range between 0.9 and 1.0 correspond to rods, and each data point represents a single counted cell. Median of each condition denoted by line. Statistical analysis performed via nested 1-way ANOVA with multiple comparisons to the “No additive” condition. Ns is p>0.05, * is p<0.05, ** is p<0.01, and **** is p<0.0001.

Cell shape also varies in *Hfx. volcanii* motility halos that form on soft agar plates. Considering that motile cells at the edge of the halos are rods and cells in the dense center are disks^38^, we hypothesized that CM would render cells non-motile. Indeed, stab-inoculating *Hfx. volcanii* into motility agar containing CM resulted in severe motility defects: no motility was observed in plates containing 50% CM (**Fig. S2**) and only limited non-uniform motility patterns started forming in plates with 25% CM (**Fig. 2**). Even at 10% CM motility halos are non-uniform (**Fig. S2**). These results are consistent with CM promoting the formation of non-motile disks.

**Figure 2.**
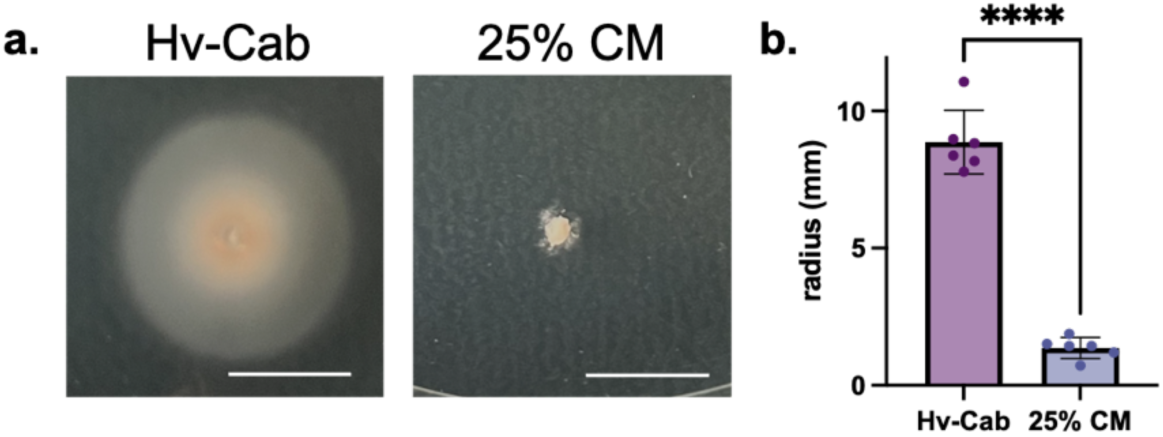
CM incorporated into soft agar plates inhibits swimming motility. Motility halos of *Hfx. volcanii* stab-inoculated onto soft agar plates of Hv-Cab (left) and Hv-Cab supplemented with 25% CM (right). Scale bar denotes 1 cm **(a)**. Quantification of motility halo radius in millimeters of six biological replicates. Error bars depict mean with standard deviation. Statistical significance determined with unpaired parametric two- tailed t-test. **** is p<0.0001 **(b)**.

### Identification of mutant strains unable to respond to CM

Two knockout strains Δ*ddfA* and Δ*cirA* were previously shown to be hypermotile and persistently rod-shaped^39,40^. To determine whether the hypermotility was a result of their potential inability to respond to a density-dependent signal in the surrounding medium, we tested their swimming motility in the presence of CM. Indeed, both Δ*ddfA* and Δ*cirA* could form uniform motility halos on soft agar plates supplemented with CM (25% v/v) (**Fig. 3a** **top, 3b**). Both strains also display rod-shaped cells in early log phase when grown in liquid cultures containing 1% CM (**Fig. 3a** **bottom, 3c**). It should be noted that CM from both Δ*ddfA* and Δ*cirA* strains did induce disk formation of wild-type *Hfx. volcanii* (**Fig. S3**). These results strongly suggest the involvement of DdfA and CirA in the recognition and/or response pathways of *Hfx. volcanii* QS.

**Figure 3.**
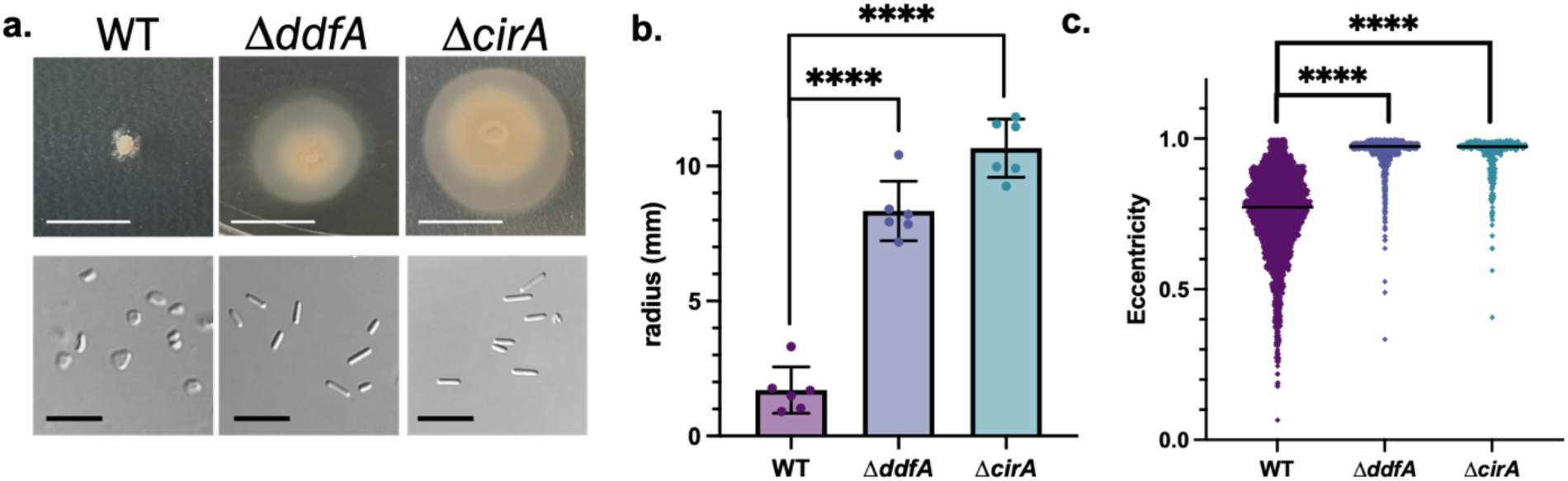
Δ*ddfA* and Δ*cirA* form motile rod-shaped cells in the presence of CM. Motility halos of wild type (WT), Δ*ddfA*, and Δ*cirA* on soft agar plates supplemented with 25% CM (v/v). Scale bar denotes 1 cm **(a, top)**. DIC microscopy images of WT, Δ*ddfA*, and Δ*cirA* cells grown to early log in cultures containing 1% CM. Scale bar denotes 10 μm **(a, bottom)**. Quantification of six replicate motility halo radii in millimeters. Error bars depict mean with standard deviation. Statistical significance determined with ordinary 1-way ANOVA with multiple comparisons to the WT condition **(b)**. Cell shape quantification of three biological replicates of WT, Δ*ddfA*, and Δ*cirA* grown to OD_600_ 0.05 containing 1% CM as described in Figure 1. Median of each condition denoted by line. Statistical analysis performed via nested 1-way ANOVA with multiple comparisons to the WT. **** p-value <0.0001 **(c)**.

### Identification of additional putative components involved in QS recognition and response pathways using quantitative proteomics

Quantitative proteomics workflows optimized for *Hfx. volcanii* have been invaluable for uncovering key components of archaeal cellular processes, such as cell shape^41,42^. Previously, proteomic changes between *Hfx. volcanii* cells at early and late log were analyzed^39^. Since those conditions correspond to low and high cell density, respectively, proteins involved in QS mediated behaviors may have significant abundance differences in these conditions, and we therefore revisited these results. Of the 2,136 proteins identified and quantified for the two conditions, 640 correspond to proteins with a significantly higher abundance in late log (log_2_ fold change > 0), and 423 correspond to proteins with a significantly lower abundance in late log (log_2_ fold change < 0, **Fig. 4a, Table S1**). This large number of proteins with differing abundances likely corresponds not only to proteins involved in a QS-related signal-transduction pathway, but also to proteins related to metabolic processes and other cellular functions that differ between early and late log growth conditions. To narrow down the list of potential proteins specifically involved in *Hfx. volcanii* QS, a similar proteomic comparison was conducted with early-log cultures grown with and without 1% CM. While the total number of identified proteins was similar (2,135 proteins), the number of proteins with significant differential abundance was strongly reduced with 171 and 65 proteins exhibiting significantly higher and lower abundance, respectively, in the presence of CM (**Fig. 4b, Table S2**). These two proteomic comparisons identified 194 proteins in common, potentially corresponding to the proteins specifically involved in the QS response pathway (**Fig. 4c, Table S3**).

**Figure 4.**
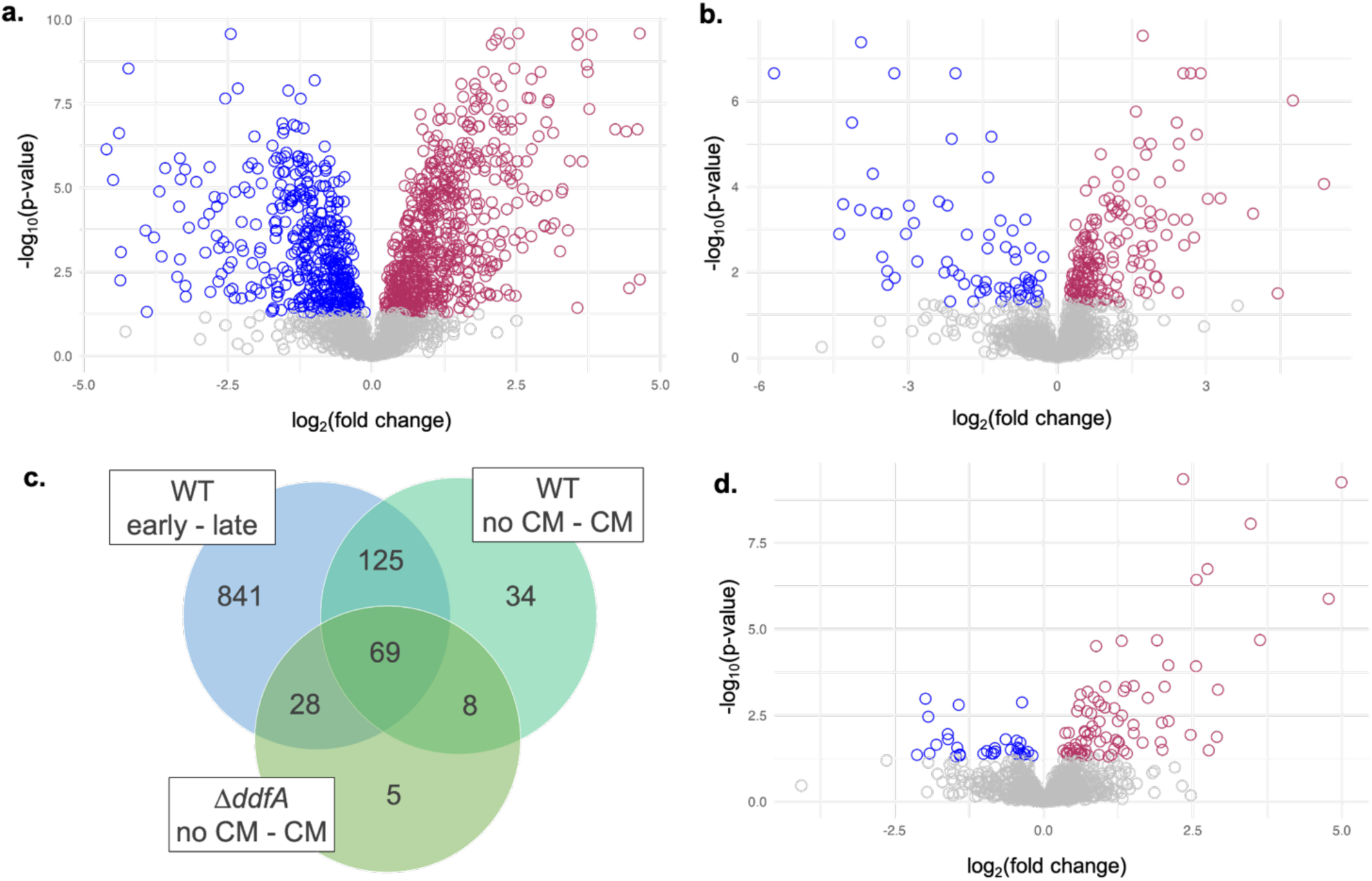
Quantitative proteomics analyses indicate proteins with altered abundances in the presence of CM. Volcano plots depicting log2 fold changes and their corresponding -log10 p-values of protein abundances compared wild type between early and late log **(a)**, wild type early log regular medium to 1% CM **(b)**, and Δ*ddfA* early log regular medium to 1% CM **(d)**. Non-significant (p-value ≥ 0.05) protein changes correspond to gray circles; significant (p-value < 0.05) and positive protein fold changes correspond to maroon circles (higher abundance in second condition); significant (p-value < 0.05) and negative protein fold changes correspond to blue circles (higher abundance in first condition). Raw values can be found in the supplementary material. Interactive html files of the volcano plots are available at 10.5281/zenodo.14510577 **(a,b,d)**. Venn diagram of the overlap of proteins with significant differential abundance in the three proteomics comparisons (wild type early to late log, wild type no CM to 1% CM, and Δ*ddfA* no CM to 1% CM). A list of proteins represented in each overlapping region can be found in Table S3 **(c)**.

Consistent with our previous study that focused on cell shape^39^, the comparison with 1% CM also identified differentially abundant proteins involved in shape: proteins important for rods (CetZ1, RdfA, and Sph3) were reduced in 1% CM and proteins important for disks (VolA and CetZ5) were increased. Similarly, in line with the decreased swimming motility of cells in the presence of CM, proteins involved in the formation and function of the flagella analog, archaella, (ArlA1 and ArlG) as well as chemotaxis (CheC, CheY, CheA, CheW1) were at lower abundance in 1% CM. Pilus production was also affected; PilA2 abundance increased while PilB1 and PibD abundance decreased in the presence of CM. Since the adhesion pilins PilA1-6 rely primarily on the PilB3/C3 complex for assembly^43^, it is likely that the PilB1/C1 complex is used for other specialized functions^44^, highlighting the potential for different pili functions in the presence or absence of CM.

Notably, CirA abundance was increased in cells grown with CM, supporting our hypothesis that CirA plays a role in recognition of and/or response to QS signals in the CM. Additionally, CirA paralogs CirB and CirD also had differential abundances but in opposite directions: CirB abundance increased in 1% CM but CirD abundance strongly decreased. Strikingly, almost all the proteins in one of the *Hfx. volcanii N*-glycosylation pathways (Agl5, Agl6, Agl7, Agl8, Agl9, Agl10, Agl11, Agl12, Agl14, and Agl15) had an increased abundance in 1% CM. Consistent with this observation, a higher percentage of glycoproteins^45^ was observed as differentially abundant in 1% CM than in the total proteins identified by mass spectrometry as well as in the proteins of differential abundance in the early log to late log comparison (**Fig. S4a**).

Not only did the proteins involved in shape and motility show significant differential abundances, the proteins important for motile rods were the ones that decrease in abundance the most (log_2_ fold change < -3): RdfA, Sph3, ArlA1, ArlG, CheY, and CheW1, signifying their importance in the QS response to CM. However, many of the other proteins that have similar significant abundance changes have unknown functions or are poorly characterized. The additional proteins that also showed a decreased abundance corresponding to log_2_ fold change < -3 included PilB1, CirD, PaaC (1,2-phenylacetyl-CoA epoxidase subunit C), HVO_1902 (conserved hypothetical protein), HVO_2074 (probably secreted glycoprotein), HVO_A0590 (conserved hypothetical protein), and HVO_B0095 (UspA domain protein). In the opposite direction, the proteins that showed a log_2_ fold change greater than 3 were BasB (chemotactic signal transduction system periplasmic substrate-binding protein), LivJ1 (ABC-type transport system periplasmic substrate-binding protein), IucA (siderophore biosynthesis protein), HVO_A0133 (conserved hypothetical protein), HVO_2836 (sensor/bat box HTH-10 family transcription regulator), and HVO_A0350 (conserved hypothetical protein). Among this set, BasB and HVO_2836 are proteins that could be potentially involved in signal transduction pathways in response to signals in the CM.

Since two component systems in Gram-positive bacteria have been shown to be involved in QS^16^, we also looked for proteins annotated as potential two-component systems among the significant differentially abundant proteins. We searched for the proteins annotated as histidine kinases as well as response regulators that have the key phosphorylation receiver domain (Interpro: IPR001789). Indeed, HVO_A0550 (receiver/sensor box histidine kinase) showed increased abundance, while HVO_1356 (sensor box histidine kinase) and HVO_1271 (receiver box response regulator) were less abundant in 1% CM. Along with HVO_1356, the nearby small CPxCG-related zinc finger protein HVO_1359 is also differentially abundant in the presence of 1% CM, showing an increased abundance. These two proteins are in the vicinity of HVO_1357, a protein with unknown function previously linked to hypermotility^46^.

Because Δ*ddfA* remains as motile rods in response to CM, a proteomic comparison between Δ*ddfA* with and without CM was also conducted to determine the effect of CM apart from shape and motility transitions. Of the 2,122 total proteins identified between Δ*ddfA* cultures at early log in regular medium versus 1% CM, 79 corresponded to proteins with significantly higher abundance in the presence of CM and 31 corresponded to proteins with significantly lower abundance (**Fig. 4d, Table S4**). Consistent with the inability for Δ*ddfA* to transition to disks, the comparison between Δ*ddfA* with and without CM did not identify previously recognized shape and motility proteins (CetZ1, RdfA, ArlG, CheC, CheY, CheA, CheW1) as having differential abundances, while other shape and motility proteins (Sph3, VolA, CetZ5, ArlA1) showed a smaller log_2_ fold change than for the wild-type comparison with CM. 69 proteins w ere identified in the intersection of the three proteomic comparisons; this set of proteins may correspond to proteins involved in *Hfx. volcanii* QS independent of DdfA (**Fig. 4c, Table S3**). Assuming that DdfA plays an essential role in the QS response pathway, these proteins could thus be involved in steps preceding DdfA or in branches that are parallel to DdfA. Pilus production was still affected in Δ*ddfA* with 1% CM: the pilin proteins PilA1 and PilA2 showed an increase in abundance while PilB1 showed a decrease. While CirA and CirB no longer exhibited significant differential abundance, CirD still had a decreased abundance in Δ*ddfA* with CM. Many proteins of the Agl15- dependent *N*-glycosylation pathway (Agl5, Agl6, Agl7, Agl8, Agl11, Agl12, Agl14) also were differentially abundant. While most of the Agl proteins were more abundant in 1% CM, similar to the comparison in wild type, Agl6 and Agl8 showed a decrease in abundance for Δ*ddfA*. Both Agl6 and Agl8 are involved in early steps of the glycan biosynthesis pathway, potentially pointing to a differential regulation of different parts of this pathway. Interestingly, a higher percentage of glycoproteins^45^ among differentially abundant proteins was found in the proteomic comparison of Δ*ddfA* with and without CM than in any of the other comparisons (**Fig. S4a**).

To get an overview of functional categories for proteins that are potentially involved in QS, we also assessed whether any arCOG categories^47^ are over- or underrepresented in the three sets of significantly differentially abundant proteins in our analyses (**Fig. S4c**). As a reference, the distribution of arCOG categories for all the proteins identified and quantified in the mass spectrometry (MS) data was used. Consistent with *Hfx. volcanii* shape and motility transitions as a result of QS, the categories significantly overrepresented in the comparisons with CM were categories M (cell wall/membrane/envelope biogenesis) and N (motility). Additionally, for the wild-type comparison with and without CM, category K (transcription) was underrepresented.

### The disk forming signal is a small molecule

After determining the cell physiological and proteomic responses to the addition of CM, we set out to gain insight into the molecular nature of the putative disk forming signal (DFS) in the CM. DFS is stable at high temperature and pressure, as autoclaving CM did not diminish the disk forming effect on early log *Hfx. volcanii* cultures (**Fig. 5a**). CM was also passed through a 3 kDa centrifugal filter to determine the relative size of DFS. While adding flowthrough to fresh *Hfx. volcanii* culture did not induce disks, washing the filter with Tris buffer, and then adding the resulting eluate to *Hfx. volcanii* culture did result in disk shaped cells (**Fig. 5a**), signifying that DFS is 3 kDa or smaller in size. Serendipitously, the Tris buffer elution of DFS from the filter also served as a buffer exchange, significantly reducing the salt concentration of the DFS-containing eluate.

**Figure 5.**
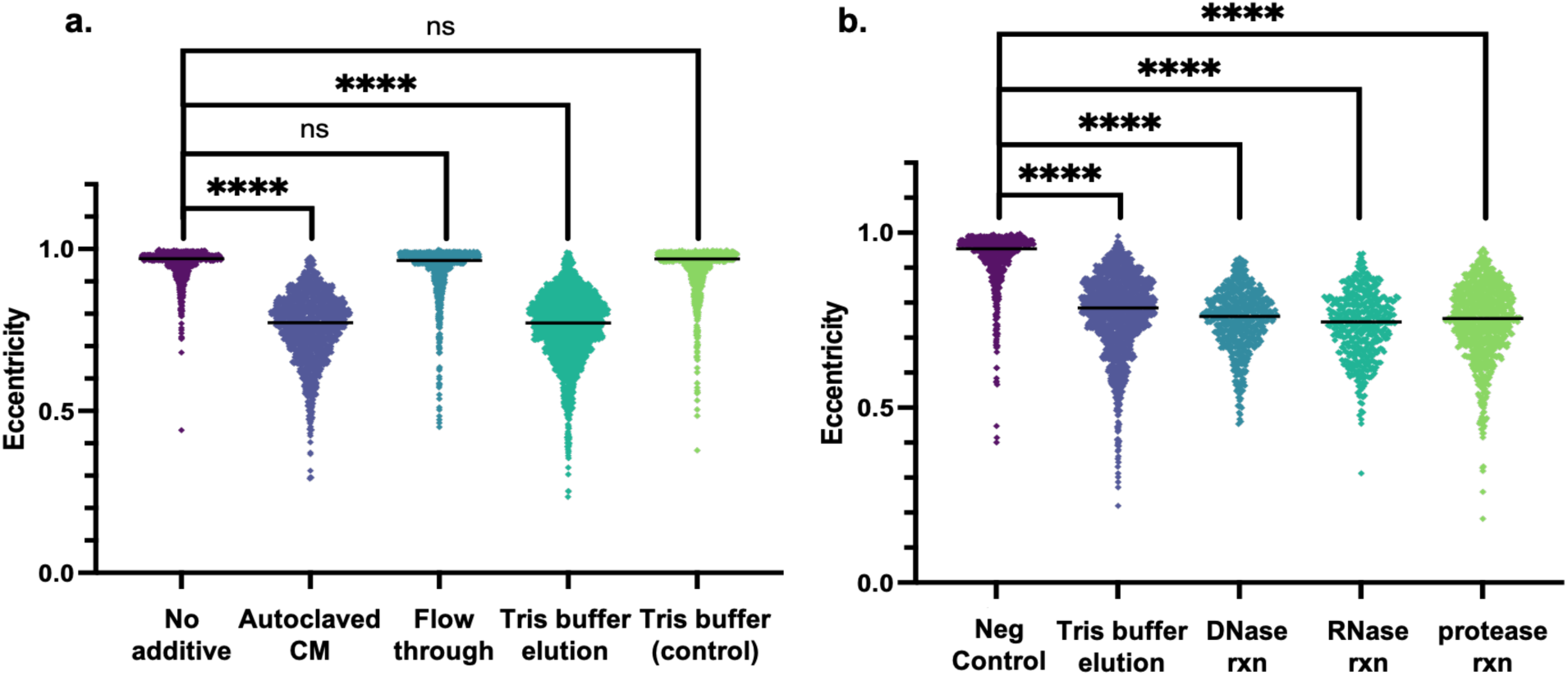
DFS can be extracted from CM and is unaffected by DNase, RNase, or Proteinase K. Cell shape quantification of three biological replicates of cultures containing final concentration of 1% of each of the following: autoclaved CM, flowthrough of CM passing through a 3 kDa centrifugal filter, elution from the filter using Tris buffer, and the Tris buffer as control **(a)**, as well as eluates treated with RQ1 DNase, RNase A and Proteinase K. “Neg control” condition includes 5 µL RQ1 DNase 10X Reaction Buffer, 5 µL of 50% glycerol, and 5 µL RQ1 DNase Stop Solution to ensure that these additional components do not induce disk formation **(b)**. Cell shape quantification as described in Figure 1. Line denotes median value. Statistical analysis performed via nested 1-way ANOVA with multiple comparisons to the respective “No additive” or “Neg control” condition. Ns is p>0.05 and **** is p<0.0001.

This low-salt DFS eluate could hence be used to test the stability of DFS against various commercially available enzymes. Treatment of eluted DFS with RQ1 DNase, RNase A, and Proteinase K did not disrupt disk formation (**Fig. 5b**), signifying that DFS is not a DNA, RNA, or protein molecule. These results strongly suggest that DFS is a small molecule.

### Nucleotide-based secondary messengers and diketopiperazine molecules do not induce disk formation

3’5’-cGMP, 3’5’-cAMP, and Ap_4_A are three nucleotide-based secondary messengers that each play roles in bacterial virulence and biofilm formation^48–50^. Since virulence and biofilm formation are commonly linked with QS systems^51,52^, and these three secondary messengers have been recovered from *Hfx. volcanii* cell pellets^53^, we explored the potential for one of these small molecules to promote the *Hfx. volcanii* QS response. We observed that applying 3’5’-cGMP, 3’5’-cAMP, or Ap4A to fresh *Hfx. volcanii* cultures did not lead to increased disk formation at early log (**Fig. S5**).

Next, we attempted the purification of DFS beginning with an extraction of CM with ethyl acetate, resulting in an organic extract and an aqueous layer. Addition of the dried organic extract to fresh *Hfx. volcanii* cultures promoted disk formation at early log, whereas addition of the remaining aqueous layer did not (**Fig. S6**). Further fractionation of the organic extract was conducted via normal phase silica gel column chromatography by eluting with solvents of increasing polarity, collecting and concentrating these fractions for bioactivity testing. Liquid chromatography tandem mass spectrometry (LC-MS/MS) was conducted on the crude organic extract as well as the fractions obtained from the column chromatography, and the resulting data was dereplicated through molecular networking (**Fig. S7**).

Among the enriched masses, we identified diketopiperazine (DKP) molecules with fragmentation patterns that matched cyclo(prolyl-tyrosine), cyclo(prolyl-valine), cyclo(prolyl- phenylalanine) and cyclo(prolyl-leucine) [or cyclo(prolyl-isoleucine), since leucine and isoleucine are isobaric amino acids] as components of CM. Because DKP molecules isolated from *Htr. hispanica* supernatant had been shown to activate an *Agrobacterium tumefaciens* NTL4 bioreporter for QS^28^, we hypothesized that the *Hfx. volcanii* DKPs play a role in archaeal QS. We show that, using the *A. tumefaciens* KYC55 (pJZ410)(pJZ384)(pJZ372) bioreporter^54^, *Hfx. volcanii* CM did indeed elicit a response (**Fig. 6a**). Thus, we hypothesized that the DKPs could be the molecules that induce the *Hfx. volcanii* shape transition. 18 of the 20 enantiomers of these identified DKPs were synthesized (we decided not to synthesize cyclo-(L-Pro-D-Ile) and cyclo-(D- Pro-D-Ile) due to the prohibitive cost of D-Ile, see supplemental material). However, none of the synthesized DKPs were found to induce disk formation in *Hfx. volcanii* (**Fig 6b**). These results highlight the potential for DFS to be a small molecule that has not been previously associated with QS.

**Figure 6.**
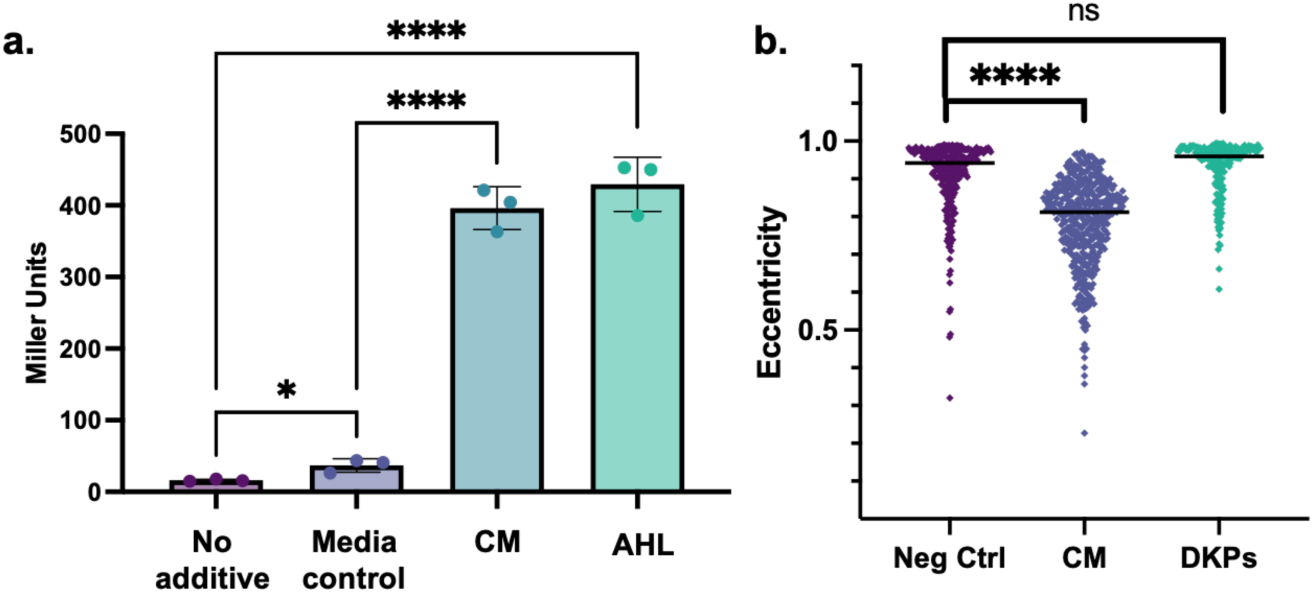
*Hfx. volcanii* CM activates *A. tumefaciens* bioreporter strain similar to AHL, but DKPs do not induce *Hfx. volcanii* disk formation. Miller Units of cleavage of ortho-Nitrophenyl-β- galactoside (ONPG) were measured in the bioreporter strain *A. tumefaciens* KYC55 (pJZ410)(pJZ384)(pJZ372)^53^. “No additive” condition corresponds to basal output of the bioreporter strain, “Media ctrl” condition corresponds to the addition of 200 µL Hv-Cab extract, “CM” condition corresponds to 200 µL of CM extract, and “AHL” condition corresponds to final concentration 2.5 ng/µL of N- 3-oxo-pentanoyl-L-Homoserine lactone. Statistical analysis performed via unpaired parametric two-tailed t-test **(a)**. Cell shape quantification of three biological replicates comparing no additive and 1% CM to the addition of DKPs pooled together for final concentration 0.2 µg/mL. Because DKPs were solubilized in ethanol, similar amounts of ethanol (10 µL) were added to the “Neg Ctrl” and “CM” cultures. Cell shape quantification as described in Figure 1. Line denotes median value. Statistical analysis performed via nested 1-way ANOVA with multiple comparisons to the respective “No additive” or “Neg control” condition **(b)**. Ns is p>0.05, * is p<0.05, and **** is p<0.0001.

## Discussion

In this interdisciplinary study, we developed a system to learn about archaeal QS using the model halophile *Hfx. volcanii*, combining microbiology with proteomics and analytical chemistry. Our results demonstrate that *Hfx. volcanii* secretes a QS signal (DFS) into the extracellular environment that regulates the rod-to-disk transition (**Fig. 1**). We show that DFS is a population-density dependent signal because the addition of increasing amounts of CM to fresh cultures as well as higher OD_600_ of culture from which CM is prepared leads to a higher proportion of disk-shaped cells at early log (**Fig. 1b, 1c**). In addition to the QS-mediated morphology transition, we show that the addition of CM to motility agar can arrest *Hfx. volcanii* motility (**Fig. 2**). A larger volume of CM is necessary for visualizing the motility cessation phenotype than for the shape transition phenotype (**Fig. S2**), possibly due to agar sequestration of DFS or shaking during liquid culture incubation. Thus, in this study we establish two distinct, easily observable and reproducible QS-mediated phenotypes in *Hfx. volcanii.* Furthermore, we demonstrate that DFS is a small molecule that can be enriched via a centrifugal filter (**Fig. 5**) and can be extracted by ethyl acetate (**Fig S5**), similar to other QS signals^9^.

Prior to our study, no proteins involved in the archaeal signaling pathways in response to QS signals had been identified. Using our model system, we were able to identify two existing mutants, Δ*ddfA* and Δ*cirA*, that retained the rod shape and their ability to swim in the presence of CM (**Fig. 3**). In fact, the previously shown hypermotility phenotypes of Δ*ddfA* and Δ*cirA*^39,40^ may be caused by their lack of DFS signal recognition and response. While little is known about the function of DdfA in disk formation^39^, our previous work has shown that CirA plays a role in transcriptional regulation of the genes encoding archaella biosynthesis: without CirA, downregulation of the *arl* genes *arlA1*, *arlI*, and *arlJ* does not occur^40^. Since we discovered *ΔddfA* and Δ*cirA* as, to the best of our knowledge, the first mutants that do not display the *Hfx. volcanii* QS response, we can now expand our understanding of the roles of DdfA and CirA in QS.

Subsequent quantitative proteomics of wild-type *Hfx. volcanii* grown with and without CM revealed significant differential abundance of 236 proteins, including significant abundance decreases of proteins important for motile rods and increases in proteins important for disks in the presence of CM (**Fig. 4b, Table S2**). We also observed an increase in CirA and proteins involved in *N-*glycosylation. Previous data suggest glycosylation may play a role in motility and shape regulation, and colonies of Δ*agl15* look similar to Δ*cirA*: smaller and darker than wild-type colonies^40,45^. Similar to previous studies^39,41^, DdfA was not identified in our MS analyses, however, further insight into its role in CM-dependent regulation was obtained from quantitative proteomics of the Δ*ddfA* strain in the presence and absence of CM. Although the morphology and motility phenotype of Δ*ddfA* suggests that this disk-defective mutant lacks certain responses to DFS, the identification of 110 proteins with significant differential abundance in Δ*ddfA* when grown in 1% CM suggests DdfA-independent CM responses (**Fig. 4d, Table S4**). Because 77 proteins overlap in the “wild type, no CM to CM” and “Δ*ddfA*, no CM to CM” comparisons (**Fig. 4c**), it is likely that those proteins are involved in the DFS response pathway independent of the role of DdfA or are responding to other components of CM in a DdfA-independent way. A subset of proteins that increases or decreases in this manner is depicted in **Fig. 7**. Similarly, the set of 159 proteins that change in abundance in the “wild type, no CM to CM” comparison but not in the “Δ*ddfA*, no CM to CM” comparison provide us with a set of proteins that are regulated in a DdfA-dependent manner in response to CM. This set includes CirA, CetZ1, ArlG, chemotaxis proteins, certain glycosylation pathway proteins, and potential proteins in two-component signaling. At the moment it is not clear whether DdfA directly responds to DFS or whether DdfA is indirectly affected by DFS. However, these quantitative proteomics results show the importance of QS in archaeal regulation, provide an invaluable dataset of proteins involved in QS responses, and give us first insights into the QS-mediated regulatory network in *Hfx. volcanii* (**Fig. 7**).

**Figure 7.**
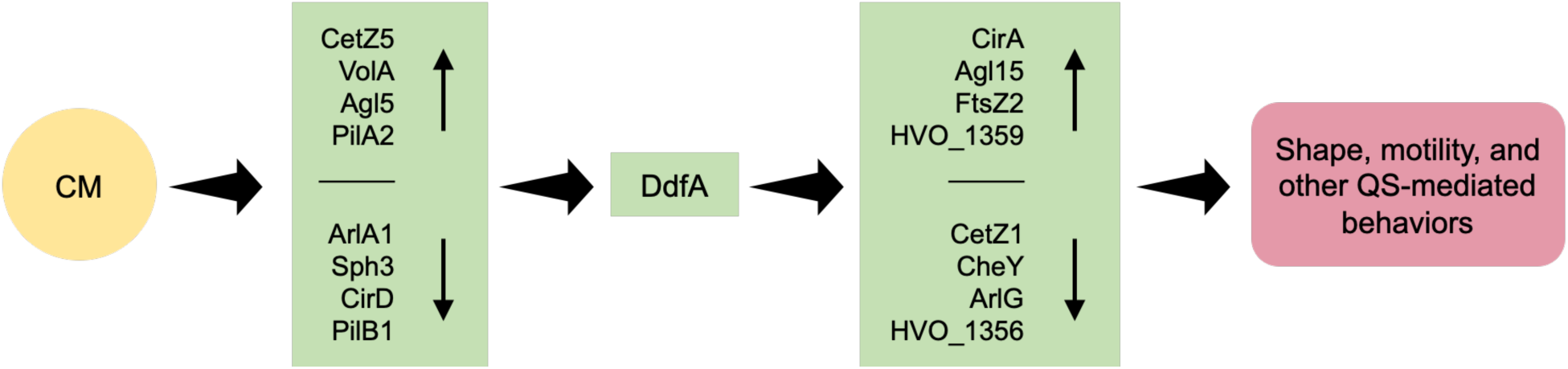
**The initial roadmap for the QS-mediated regulatory network in *Hfx. volcanii***. A subset of proteins discussed in this study is depicted as increasing or decreasing in abundance (vertical arrows) as a result of direct or indirect regulation (horizontal arrows), resulting in QS- mediated behaviors.

Finally, our data reveal not only the importance of QS in archaea but also provide a strong foundation for further studies into archaeal QS using the model archaeon *Hfx. volcanii*. We showed that DFS is a stable small molecule (**Fig. 5**), but it also appears that DFS is highly potent at low concentrations (**Fig. 1b**), rendering chemical isolation and purification difficult. We also cannot exclude that more than one molecule is responsible for the QS-mediated phenotypes. Because there is a possibility that DFS is synthesized via an autoregulatory pathway, similar to AHL in the LuxI/LuxR system^15^, the proteins involved in DFS biosynthesis may be found in our proteomics datasets and could shed light on the chemical structure of this QS signal. Therefore, future studies using multi-pronged approaches that combine chemical purification strategies and screens for mutants unable to produce DFS, along with the proteomic results obtained from this study, will aid in delineating the *Hfx. volcanii* DFS pathway. These interdisciplinary approaches will not only advance our understanding of how *Hfx. volcanii* communicates with itself, but also help us to learn more about interspecies and inter-domain interaction.

## Methods

### Growth conditions

The strains used in this study are listed in Table S5. *Hfx. volcanii* strains were grown at 45°C in liquid (orbital shaker at 250 rpm, 1-in orbital diameter) or on solid agar (1.5% w/v) in semi-defined Casamino Acids (Hv-Cab) medium^37^, supplemented with tryptophan (+Trp) (Fisher Scientific) and uracil (+Ura) (Sigma) at a final concentration of 50 µg mL^−1^. Solid agar plates are placed within sealable bags during incubation to prevent the plates from drying out.

Preparation of conditioned medium (CM) and CM-agar plates *Hfx. volcanii* H53^55^ was inoculated from 1 colony into 10 mL of Hv-Cab and incubated shaking overnight. This 10 mL starter culture was added to 1 L of Hv-Cab. OD_600_ of the large culture was monitored by removing aliquots and checking using a spectrophotometer (Thermo Spectronic 20+). Large culture was grown until OD_600_ 1.5 (2-3 days) and then pelleted (Beckman Coulter Avanti J-25I, JA-10 rotor, 8,000 xg, 30 min). Supernatant was sterile filtered (Millipore Stericup, 0.22 µm PVDF membrane) and stored at -20°C. Soft agar plates for motility assays were prepared as 0.35% w/v agar of either only Hv-Cab medium or Hv-Cab medium supplemented with CM for final CM concentration of 25%.

### Cell shape assay

Glass culture tubes were prepared with 5 mL Hv-Cab medium corresponding to the number of experimental conditions and replicates. For each experiment, additives (i.e. CM, DKPs, etc) were introduced to the medium before inoculation. *Hfx. volcanii* was inoculated from a single colony into 1 mL of Hv-Cab liquid as starter inoculum and vortexed thoroughly for homogenization. OD_600_ of 200 µL of the starter inoculum was determined (BioTek PowerWave XS2, Gen5 1.11). The starter inoculum was used to inoculate 5 mL cultures for 1 replicate so that the cultures would uniformly reach OD_600_ 0.05 the following day (i.e. 50 µL of OD_600_ 0.055 of starter inoculum in 5 mL cultures incubated at 5:30 pm will reach OD_600_ around 11:00 the following day). Once the cultures reached OD_600_ 0.05, 1 mL of the culture was pelleted at 4,900 xg for 6 minutes. Supernatant was aspirated to leave ∼5 μL liquid remaining with the pellet (200 times concentrated). Cells in the resulting pellet were used for live-cell imaging.

### Live-cell imaging and analysis

1.5 µL of resuspended cell pellets prepared for imaging was placed on glass slides (Fisher Scientific) and a glass coverslip (Globe Scientific Inc.) was placed on top. The coverslip was gently pushed on the glass slide until the coverslip no longer moved easily, to maintain a single layer of cells to visualize. Slides were visualized using a Leica Dmi8 inverted microscope attached to a Leica DFC9000 GT camera with Leica Application Suite X (v. 3.6.0.20104) software. Differential interference contrast (DIC) as well as Brightfield images were captured at 100x magnification.

Brightfield images used for quantification were processed first with a background subtraction method on Fiji (ImageJ, version 2.9.0/1.54f) and then cells were quantified using CellProfiler (version 4.1.3)^56,57^. The Fiji macro and CellProfiler pipeline used to analyze and quantify the images can be found at https://doi.org/10.5281/zenodo.14510577. Cell-specific data for each image set were exported. The eccentricity parameter was used for determining the shape differences and graphed on GraphPad Prism version 9.3.1 and 10.0.2 for macOS. Statistical significance of eccentricity comparisons between conditions were assessed with a nested 1-way ANOVA with multiple comparisons to the negative control condition for comparisons across multiple datasets using GraphPad Prism versions 9.3.1 and 10.2.1 for macOS (GraphPad Software, San Diego, California USA, www.graphpad.com). Frequency plots using eccentricity values were graphed using R Studio (version 2024.04.2+764) and ggplot2 (version 3.5.1). Script can be found on https://doi.org/10.5281/zenodo.14510577.

### Motility assays and motility halo quantification

A toothpick was used to pick a single colony and stab-inoculate 0.35% w/v agar plates. Plates were incubated at 45 °C upright in a plastic box with added wet paper towels to maintain humidity. Paper towels were changed daily. Motility assay plates were removed from the 45 °C incubator after two days and imaged after one day of room temperature incubation. Incubation at room temperature for one day allows darker pigmentation to be produced by the cells, making the halos easier to observe and image.

For halo radius quantification, images were uploaded to Fiji (ImageJ, version 2.9.0/1.54f), and the scale was set based on plate diameter. Statistical significance of halo diameters was assessed with an unpaired, two-tailed parametric t-test for a comparison between two datasets, or with an ordinary 1-way ANOVA with multiple comparisons to the negative control condition for comparisons across multiple datasets using GraphPad Prism versions 9.3.1 and 10.2.1 for macOS (GraphPad Software, San Diego, California USA, www.graphpad.com).

Label-Free quantitative proteomics of early log cultures in the presence and absence of CM Samples for quantitative proteomics analysis were prepared and analyzed as described in Schiller *et al.*^39^. Briefly, an initial inoculum for each strain was prepared by resuspending an individual colony in 1 mL liquid medium. Afterwards, the OD_600_ of each inoculum was measured and adjusted in 5 mL of liquid medium so that every strain and replicate started at the same cell density. For each strain, 3 biological replicates were prepared, using different colonies for inoculation. For each biological replicate, the same initial inoculum was used for samples in the presence and absence of 1% CM by transferring the corresponding amount of cells from the initial inoculum (without added CM) to 5 mL of liquid media with and without 1% CM, respectively. Cultures were then incubated at 45 °C under shaking conditions and sampled once they reached an OD_600_ between 0.045 and 0.050. 4 mL of cells were spun down, the supernatant was concentrated to <100 µL using centrifugal filter units (3-kDa molecular weight cutoff, Millipore), and cell pellets were frozen at -80 °C.

After cell lysis and protein extraction (as described in Schiller *et al.*^39^), 20 µg of proteins were digested with trypsin using S-trap mini spin columns (ProtiFi) following manufacturer’s instructions, as described previously^42^. Peptides were desalted using home-made stage tips, and analyzed by HPLC-MS/MS using a Dionex Ultimate 3000 RSLCnano UPLC (Thermo Scientific) linked to an Orbitrap Eclipse™ mass spectrometer (Thermo Scientific) using identical parameters as in Schiller *et al*.^39^. The corresponding gradient conditions and instrument settings are listed in Table S6.

### Bioinformatic analysis of MS results

Bioinformatic processing of MS files for the identification and quantification of peptides and proteins was performed as described previously^39^. Briefly, using the Python framework Ursgal (version 0.6.9)^58^ MS raw files were converted into mzML and MGF format with the ThermoRawFileParser (version 1.1.2)^59^ and pymzML (version 2.5.0)^60^, respectively. MSFragger (version 3.0)^61^, X!Tandem (version vengeance)^62^, and MS-GF+ (version 2019.07.03)^63^ were employed for protein database searches against the theoretical proteome of *Hfx*. *volcanii* (4,074 proteins, version 190606, June 6, 2019, https://doi.org/10.5281/zenodo.3565631), supplemented with common contaminants, and decoys using shuffled peptide sequences. Precursor mass tolerance was set to 10 ppm, and a fragment mass tolerance to 15 ppm. The following modifications were included: oxidation of M (optional), N-terminal acetylation (optional), carbamidomethylation of C (fixed). Percolator (version 3.4.0)^64^, was used for the calculation of PEPs, and results from different search engines were combined as described before^65^. PSMs with a combined PEP > 1% were removed and for spectra with multiple differing PSMs, only the PSM with the best combined PEP was accepted.

Peptide intensities were determined using FlashLFQ (version 1.1.1)^66^, with the match-between-run option activated (retention time window: 0.5 min), and were then further processed using MSstats (version 4.0)^67^. Any FlashLFQ peptide intensities with a value of ‘0’ were replaced with ‘NA’, and decoy peptides as well as contaminants were removed. The data set was then transformed into the MSstats input format using the columns ProteinName, PeptideModifiedSequence, PeptideSequence, Condition, and BioReplicate. IsotopeLabel was set to ‘L’ for all rows, and PrecursorCharge, ProductCharge, and FragmentIon were set to ‘NA’. The MSstats dataProcess() function was used with all default parameters (including logarithmic transformation and median normalization), with the exception of the model-based imputation (MBImpute was set to ‘FALSE’). After generating a contrast matrix with the desired condition comparisons, the groupComparison() function was run to obtain log_2_ fold-changes with corresponding adjusted p-values.

Functional annotations, along with COG and arCOG categories, for *Hfx. volcanii* proteins were obtained from eggNOG-mapper^68,69^ and the ftp server for arCOG indices. The arCOG categories were matched to the MSstats output of identified and quantified proteins in R Studio (version 2024.04.2+764). ‘NA’ and ‘Inf’ values were removed, along with peptides that were unable to be resolved to one protein, and then volcano plots were generated using ggplot2 (version 3.5.1) and plotly (version 4.10.4). *Hfx. volcanii* glycoprotein^45^ abundance was manually counted for each dataset and graphed using ggstatsplot (version 0.12.5)^70^. Proteins corresponding to each arCOG category were counted in R Studio (version 2024.04.2+764) and the percent of proteins identified as significantly differentially abundant in the three proteomics comparisons over the percent of all identified and quantified proteins was graphed using GraphPad Prism version 10.2.1 for macOS (GraphPad Software, San Diego, California USA, www.graphpad.com). R scripts and interactive volcano plots can be found at https://doi.org/10.5281/zenodo.14510577.

### Passing CM through 3 kDa centrifugal filter

0.5 mL of CM was added to a centrifugal filter unit (Amicon® Ultra, 0.5 mL, 3 kDa) and centrifuged at 13,000 xg for 6 minutes. 0.2 mL of CM was applied to the top of the centrifugal filter and centrifuged again at 13,000 xg for 6 minutes. The flowthrough and concentrate were removed and stored separately. 0.4 mL of 10 mM Tris-HCl (pH 8.0), 0.1 mM EDTA buffer (invitrogen) or 5 mM Tris/HCl (pH 8.5) buffer (Macherey-Nagel) was applied to the filter and centrifuged at 13,000 xg for 6 minutes. The subsequent flowthrough was collected and used for further experimentation.

### RQ1 DNase, RNase A, and Proteinase K reactions

Each reaction totaled 170 µL. 136 µL of elution in 5 mM Tris/HCl (pH 8.5) buffer (Macherey-Nagel) from the 3 kDa centrifugal filter was distributed into three separate microcentrifuge tubes labeled for the respective reaction. For the DNase reaction, 17 µL of RQ1 DNase 10X Reaction Buffer and 17 µL of RQ1 RNase-free DNase (Promega) were added. For the RNase reaction, 17 µL of 5 mM Tris/HCl (pH 8.5) buffer (Macherey-Nagel) and 17 µL of RNase A Solution (Thermo Scientific) were added. For the protease reaction, 17 µL of 5 mM Tris/HCl (pH 8.5) buffer (Macherey-Nagel) and 17 µL of Proteinase K solution (Thermo Scientific) were added. Each microcentrifuge tube reaction was incubated in a heat block at 37°C for 45 minutes. 17 µL of RQ1 DNase Stop Solution (Promega) was added to the DNase reaction and 1 µL of 200 mM AEBSF (final concentration 1 nM) was added to the protease reaction. All three reactions were heat inactivated at 65°C for 10 minutes. 50 µL of each reaction was added to triplicate fresh *Hfx. volcanii* cultures to determine if DFS was inactivated.

### Organic extraction of CM

500 mL of CM was added to a large (1L) separatory funnel and extracted with hexanes (3x 200mL), then methylene chloride (3x 200 mL), then ethyl acetate (3x 200 mL). The combined ethyl acetate extracts were concentrated to dryness using a rotary evaporator. An emulsion formed upon the addition of ethyl acetate, which required additional time for settling.

### Column chromatography of CM extract

A 10 mL column was packed with silica gel. The dried CM extract (23 mg) was dissolved in 4-5 drops of ethyl acetate and applied to the top of the column. 50 mL (5 column volumes) of each of the following solvent systems were passed through the column: 1) hexanes 2) 50% hexanes, 50% ethyl acetate 3) ethyl acetate 4) methylene chloride 5) 50% methylene chloride, 50% methanol. The eluent from each solvent system was collected separately and dried using a rotary evaporator.

### Identification of DKPs in CM

Detection of compounds in the CM extracts and column fractions was carried out using an Agilent 1290 UPLC connected to an Agilent 6545 qTOF equipped with a Dual Agilent Jet Stream Electrospray Ionization source. An Agilent InfinityLab Poroshell 120 EC-C18 (100 x 2.1 mm, 2.7 μm particle size) was equipped with a UHPLC guard column of the same packing chemistry and held at 30 °C for the separation of the extracts. The solvent system for separations consisted of solvent A (water + 0.1% formic acid) and solvent B (acetonitrile + 0.1% formic acid). The UPLC method used began with a 0.5 min hold of 95% A, then a linear gradient from 95% A to 5% A over 5.5 min. The flow rate was held constant at 0.8 mL/min. The MS method for ESI in positive mode used the following settings: gas temp, 320 °C; drying gas, 8 L/min; nebulizer, 35 psi; sheath gas, 350 °C, 11 L/min; VCap 2500 V; Nozzle Volt. 1000 V; m/z 50 to 3000; 5 spectra/s. The MS/MS method for ESI in positive mode used the same settings as the MS method, expect for the following: 2 spectra/s, collision energy calculated using CE = [5*(m/z)]/100; precursor selection of 5 masses per cycle. Data were acquired using Agilent MassHunter. Raw data were converted to the open data format .mzML^71^ using MSConvert from ProteoWizard^72^.

MS1 and MS2 features were extracted using MZmine2.53 (enabled for feature-based molecular networking) following the workflow for feature based molecular networking (FBMN) using the parameters shown in **Table S7**^73^. The detected feature list (.csv), fragmentation patterns (.mgf), and .mzML data files (listed in **Table S8** and uploaded to Zenodo https://doi.org/10.5281/zenodo.14510577) were exported and uploaded to GNPS for molecular networking and dereplication. The following description of the GNPS process parameters has been adapted from Wang *et. al.*^74^: “The FBMN module in GNPS first filters data by removing all MS/MS fragment ions within ±17 Da of the precursor m/z. The top 6 fragment ions in the ±50 Da window of each MS/MS spectra were chosen for downstream processing. The precursor m/z tolerance was set to 0.02 Da and the MS/MS m/z tolerance was set to 0.02 Da. The molecular network was generated such that edges were drawn between two nodes that shared more than 6 matched peaks in their MS/MS spectra and shared a cosine similarity score above 0.7. The resulting list of edges was filtered to contain only those where each node connected was in the top 10 list of most similar nodes for the other. The maximum size of a molecular family was set to 100, with lowest scoring edges removed from the molecular families until the molecular family size was below this threshold. The processed spectra in the network were then searched against the GNPS spectral libraries^74,75^, which were then filtered using the same parameters as the experimental input data. A positive match between the experimental and library spectra was identified by at least 6 matching peaks in the MS/MS spectra and a cosine score above 0.7.” The resulting FBMN network was downloaded as a .graphml file for visualization and annotation in Cytoscape (v. 3.10.3).

### Synthesis of DKPs

Cyclo-(L-Pro-L-Phe) and cyclo-(L-Pro-L-Tyr) were commercially available and tested directly. Commercially available substrates were used in the synthesis of the remaining DKPs for testing following known synthetic methods: N-Boc-L-Tyr, N-Boc-L-Phe, L-Pro-OBn HCl, D-Pro-OMe HCl, and all starting amino acids. See “Detailed schematics for synthesis of DKP molecules” in the supplemental material for thorough description of synthesis schemes^76–82^.

### Agrobacterium assay

The *A. tumefaciens* KYC55 (pJZ410)(pJZ384)(pJZ372) bioreporter^54^ was grown and pre-induced in accordance with Joelsson & Zhu 2006^83^. 25 mL of Hv-Cab and CM was mixed with 25 mL ethyl acetate and let settle to form two immiscible layers. Emulsions caused by the high salinity were resolved by centrifugation. The organic layer was evaporated to result in the respective Hv-Cab or CM extract. 2 µL of pre-induced *A. tumefaciens* was added to 2 mL of AT medium^83^, along with either no additive, 200 µL of 5 mg/mL each extract, or N-3-oxo-pentanoyl-L-Homoserine lactone at a final concentration of 2.5 ng/µL. Cultures incubated at 30°C (orbital shaker at 250 rpm, 1-in orbital diameter). When cultures reached OD_600_ 0.2-1.0, 200 µL of the cultures were transferred into separate microcentrifuge tubes and combined with 0.8 mL Z buffer, 10 µL 0.05% SDS, and 15 µL chloroform and vortexed. 0.1 mL of 4 mg/mL ortho-Nitrophenyl-β-galactoside (ONPG) was added to each solution and the time was recorded. When solutions turned yellow, 0.6 mL of 1 M Na_2_CO_3_ was added to stop the reaction and the elapsed time was recorded in minutes. Miller units were calculated according to Joelsson & Zhu 2006^83^.

## Data and Code Availability

The mass spectrometry proteomics data have been deposited to the ProteomeXchange Consortium via the PRIDE^84^ partner repository with the dataset identifier PXD059278. Raw data for cell shape analysis, scripts used for the analysis, as well as LC-MS/MS files (.mzML) used for molecular networking can be found on Zenodo (https://doi.org/10.5281/zenodo.14510577). Analyzed data are provided with this paper and in the supplementary documentation.

## Supporting information

Fig S1-S7, Table S5-S8, Synthesis schematics

## Acknowledgements

We thank Sarah Hegarty for her statistical expertise for the eccentricity figures, Yirui Hong for her assistance in proteomic sample preparation, and Dr. Jun (Jay) Zhu for providing the *Agrobacterium tumefaciens* bioreporter strain. We also thank the Pohlschröder Lab for helpful discussion. We gratefully acknowledge the quantitative proteomics analysis conducted by Dr. Jeremy L. Balsbaugh and Dr. Jennifer C. Liddle of the UConn Proteomics & Metabolomics Facility, a component of the Center for Open Research Resources and Equipment at the University of Connecticut. We also gratefully acknowledge Research Computing at the Rochester Institute of Technology (RIT) for providing computational resources and support that have contributed to the research results reported in this publication.

This work was supported by the National Science Foundation grant NSF-MBC2222076 and a University of Pennsylvania Research Fund grant. P.C. was additionally supported by a National Institutes of Health Training Grant T32 GM007229 “Cell and Molecular Biology” as well as a National Institutes of Health Ruth L. Kirschstein National Research Service Award 1F31AI181536-01. C.C. was supported by the National Science Foundation Graduate Research Fellowships Program. S.S. and K.W. were also supported by the College of Science at RIT.

## Notes

https://doi.org/10.5281/zenodo.14510577

### Competing Interest Statement

The authors have declared no competing interest.

