## Supplementary material for "Quorum sensing mediates morphology and motility transitions in the model archaeon *Haloferax volcanii*": Fig S1-S7, Table S5-S8, Synthesis schematics

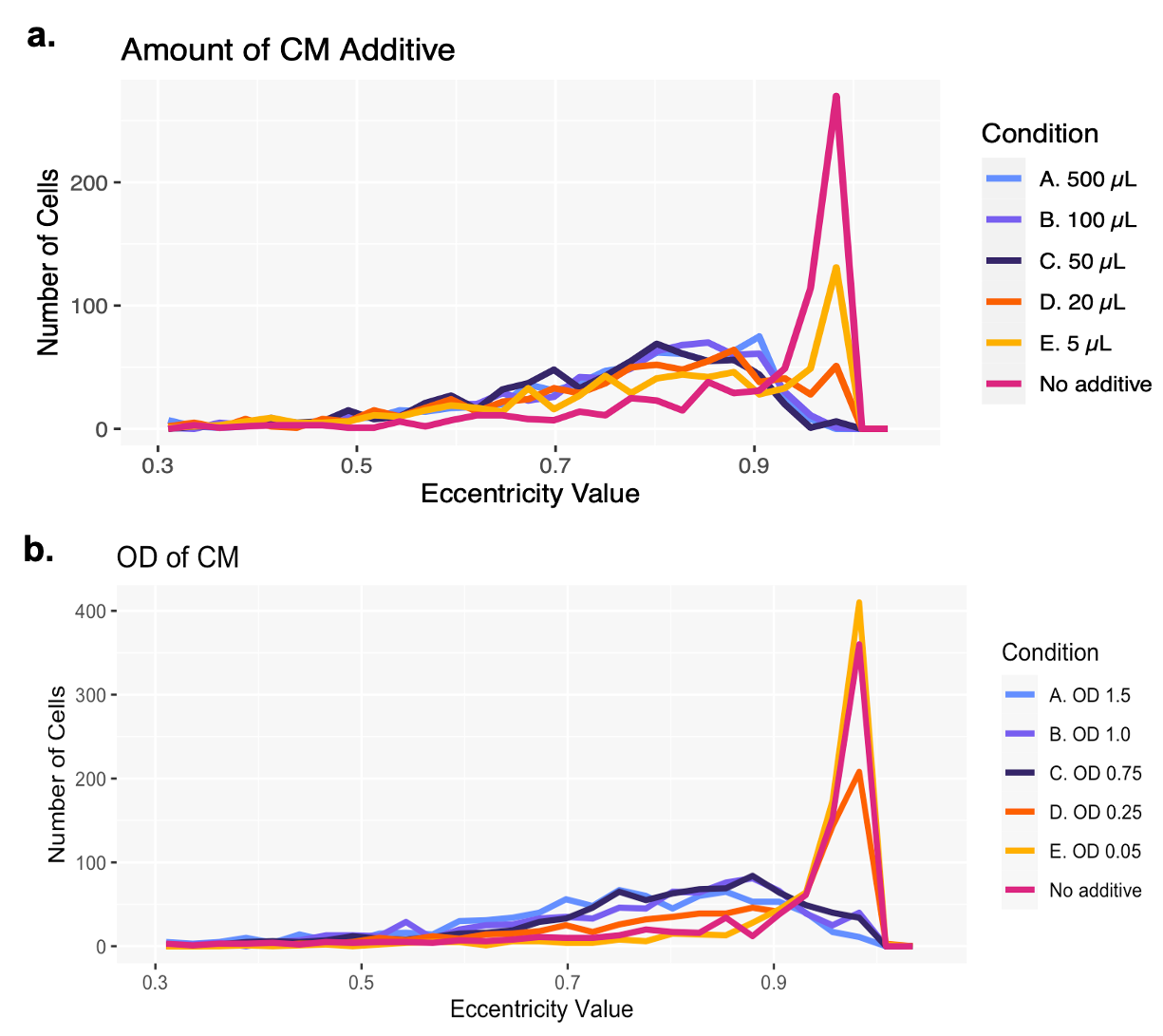

**Figure S1. Decreasing number of rod-shaped cells are observed as more CM or CM from a culture grown to a higher OD is added to fresh cultures.** Overlaid frequency plots of eccentricity measurements of *Hfx. volcanii* cells harvested at OD_600_ 0.05 from 5 mL cultures with varying volumes of added CM **(a)** and the addition of 50 µL of CM prepared from cultures at varying OD_600_ **(b)**.

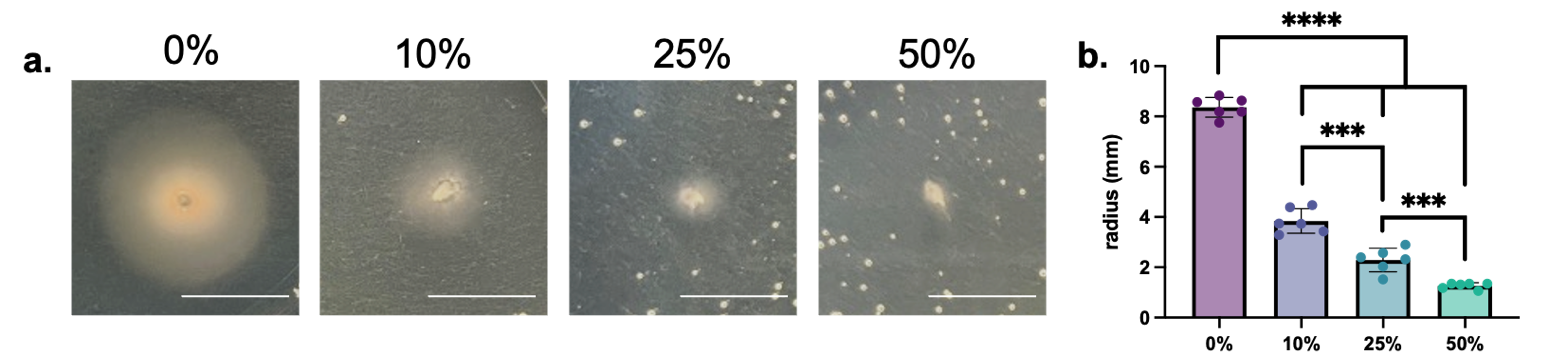

**Figure S2. Increasing percentage of CM incorporated into soft agar plates corresponds to reduction of *Hfx. volcanii* halo radius.** Motility halos of *Hfx. volcanii* cells stab-inoculated into soft agar plates of Hv-Cab supplemented with 0, 10, 25 or 50% (v/v) CM. Scale bar is 1 cm **(a)**. Quantification of motility halo radius in millimeters of six biological replicates. Error bars depict mean with standard deviation. Statistical significance determined with ordinary 1-way ANOVA with multiple comparisons to the 0% condition across all sets or unpaired parametric two-tailed t-test between two sets. *** is p<0.001, **** is p<0.0001. Bubbles sometimes appear on motility plates after incubation for unknown reasons and do not correlate to percentage of CM **(b)**.

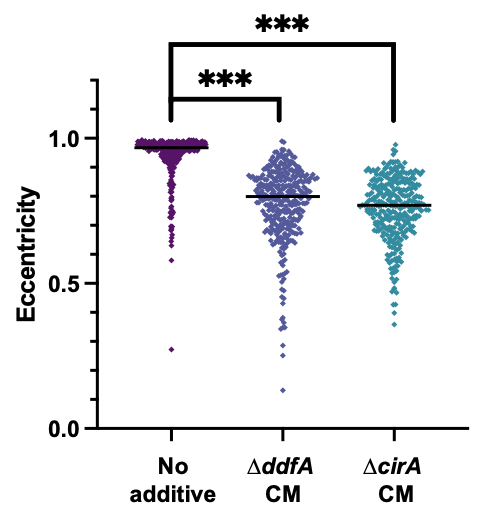

**Figure S3. CM from ∆*ddfA* and ∆*cirA* induces disk formation in wild-type *Hfx. volcanii*.** Cell shape quantification of three biological replicates of wild-type *Hfx. volcanii* cells grown to early-log in cultures containing either no additive or CM from late log cultures of ∆*ddfA* or ∆*cirA* to final concentration of 1%. Quantification as described in Figure 1. Line denotes median value. Statistical analysis performed via nested 1-way ANOVA with multiple comparisons to the “No additive” condition. *** is p<0.001.

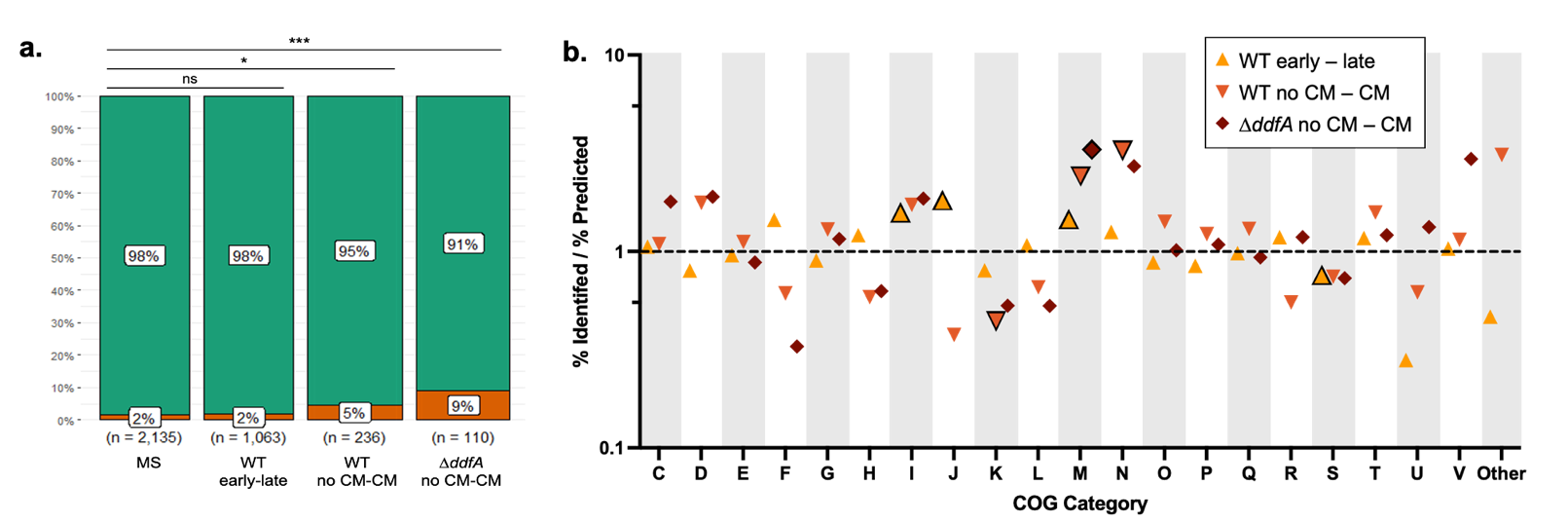

**Figure S4. Analysis of quantitative proteomics provides insight into protein abundance changes as a result of the addition of CM.** Bar plot showing the percentage of total identified and quantified proteins by mass spectrometry (MS) that are glycoproteins versus non-glycoproteins in the first bar. Subsequent bars represent the percent glycoproteins versus non-glycoproteins of the proteins that had significant differential abundance in the three proteomic comparisons. Statistical significance as determined by Fisher’s Exact test with the Bonferroni adjustment. Ns is p>0.05, * is p<0.05, and *** is p<0.001 **(a)**. For each arCOG category, the percent of proteins identified as significantly differentially abundant in the three proteomics comparisons over the percent of all identified and quantified proteins is shown. Black outlines around the data points indicate statistically significant observations as determined by Fisher’s Exact test with the Bonferroni adjustment, p≤0.05 **(b)**.

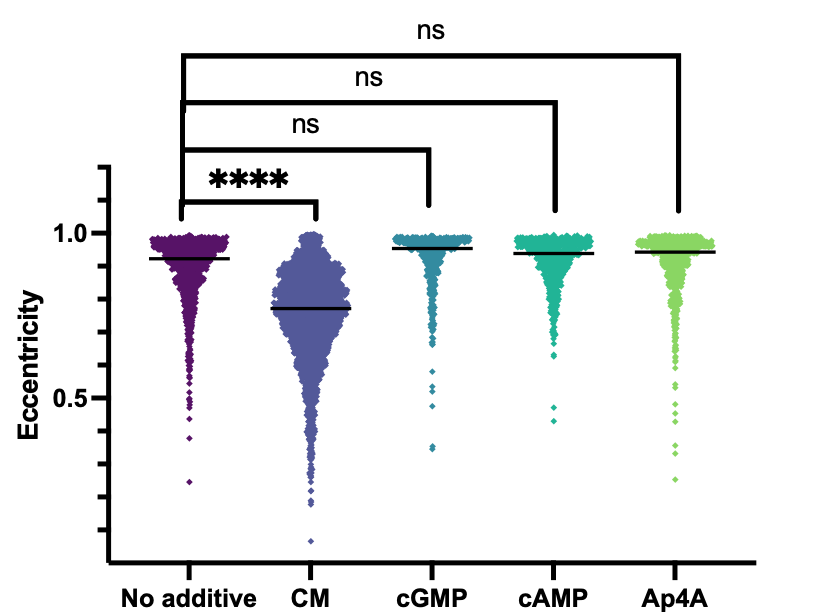

**Figure S5. Secondary messengers (3’5’-cGMP, 3’5’-cAMP, and Ap_4_A) do not promote *Hfx. volcanii* disk formation.** Cell shape quantification of three biological replicates comparing no additive and 1% CM to the addition of three secondary messengers 3’5’-cGMP, 3’5’-cAMP, and Ap_4_A each for final concentration 0.2 µg/mL. Cell shape quantification as described in Figure 1. Line denotes median value. Statistical analysis performed via nested 1-way ANOVA with multiple comparisons to the “No additive” condition. Ns is p>0.05 and **** is p<0.0001.

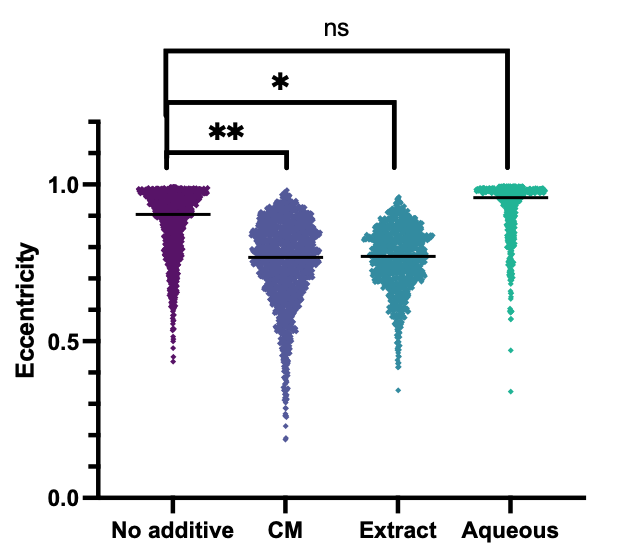

**Figure S6. Ethyl acetate can extract DFS from the CM.** Cell shape quantification of two biological replicates comparing no additive to CM, ethyl acetate crude extract of CM, and the remaining aqueous layer after extraction at final concentration of 1%. Cell shape quantification as described in Figure 1. Line denotes median value. Statistical analysis performed via nested 1-way ANOVA with multiple comparisons to the “No additive” condition. Ns is p>0.05, * is p<0.05, and ** is p<0.01.

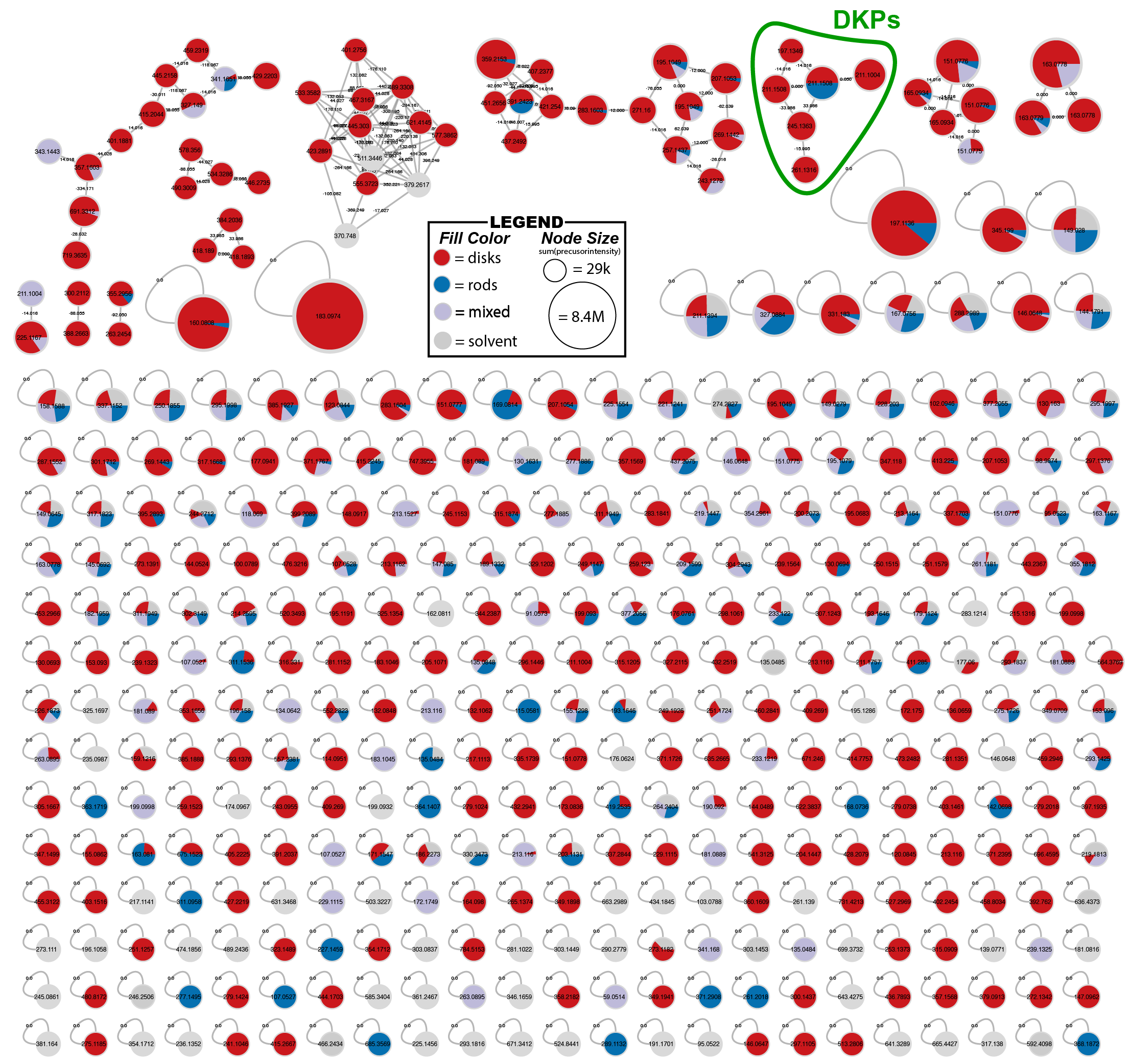

**Figure S7. Molecular network depicting the relationship between unique molecular features in the CM extract, column fractions, and solvent controls.** Individual nodes represent individual features with unique m/z and retention times. Nodes are colored in correspondence with the phenotype shown when each sample was screened for its disk-forming activity: red nodes or wedges indicate features found in disk-forming samples, blue nodes or wedges indicate features found in rod-forming samples, purple nodes or wedges indicate features found in samples that gave mixed rods and disks, and the grey nodes or wedges indicate features found in the solvent negative controls. The lines draw between nodes (edges) indicate features that share a cosine similarity score above 0.7 and more than 6 matching peaks in their MS/MS spectra. The cluster of nodes circle in green contains molecular features that were identified via de-replication in GNPS as candidate diketopiperazine molecules, based on their shared MS/MS peak patterns with GNPS library standards.

**Table S5**

List of strains and their sources used in this study

| **Name** | **Relevant characteristics** | **Source** |
| --- | --- | --- |
| *Haloferax volcanii* strains | | |
| H53 (wild type) | ∆*pyrE2* ∆*trpA* | ^59^ |
| ∆*ddfA* | ∆*pyrE2* ∆*trpA* ∆*ddfA* | ^44^ |
| ∆*cirA* | ∆*pyrE2* ∆*trpA* ∆*cirA* | ^45^ |
| *Agrobacterium tumefaciens* strain | | |
| KYC55 | pJZ410, pJZ384, pJZ372 | ^58^ |

**Table S6**

Parameters for the mass spectrometric analysis of proteomics samples.

| **Chromatography** | |
| --- | --- |
| Column | nanoEase M/Z Peptide BEH C18 column, Waters Corporation, 1.7 um particle size, 75 um x 250 mm |
| Column oven | 50°C |
| Flow rate | 300 nl/min |
| Buffer system | Buffer A: 0.1% formic acid in H_2_O |
|  | Buffer B: 0.1% formic acid in acetonitrile |
| Gradient | 1 min 3% B; increase to 17% B over 39 min; increase to 32% B over 20 min; increase to 90% B over 2 min; 9 min 90% B; decrease to 3% B over 2 min; 17 min 34% B |
| **Mass spectrometry** | |
| Ion mode | positive |
| Excluded charge states | 1, >6, unknown |
| Dynamic exclusion | 10 s |
| MS1 Resolution | 120,000 |
| MS1 Maximum injection time | 50 ms |
| MS1 AGC target | 250% (1e6) |
| MS1 Detector | Orbitrap |
| MS1 Scan range | 400-2000 *m/z* |
| MS2 Resolution | 15,000 |
| MS2 Detector | Orbitrap |
| MS2 AGC target | 200% (1e5) |
| MS2 Maximum injection time | dynamic |
| MS2 Scan range | automatic |
| MS2 Data-dependent acquisition | cycle time 3 s |
| Fragmentation | Higher-energy collisional dissociation |
| MS2 Normalized collision energy | stepped: 25, 30, 35 |

**Table S7**

MZmine2.53 Data Processing Settings

| Mass detection | MS1 | 1.0E3 |
| --- | --- | --- |
|  | MS2 | 1.0E2 |
|  | retention time | 0.5 – 5.50 min |
| Chromatogram building (ADAP) | min group size in # scans | 3 |
|  | group intensity threshold | 3.0E3 |
|  | Min highest intensity | 1.0E3 |
|  | m/z tolerance | 0.001 m/z or 5.0 ppm |
| Deconvolution (Local minimum search algorithm) | Chromatographic Threshold | 1% |
|  | Search minimum in RT range (min) | 0.2 |
|  | Minimum relative height | 2% |
|  | Minimum absolute height | 5.0E3 |
|  | Min ratio of peak top/edge | 5 |
|  | Peak Duration range (min) | 0.01 – 2.00 min |
|  | m/z center calculation | MEDIAN |
|  | m/z range for MS2 scan pairing | 0.01 Da |
|  | RT range for MS2 scan pairing | 0.1 min |
| Alignment (Join Aligner) | m/z tolerance | 0.001 m/z or 5.0 ppm |
|  | Weight for m/z | 75 |
|  | Weight for RT | 0.1 min |
|  | RT tolerance | 25 |

**Table S8**

LC-MS/MS Sample Metadata

| **filename** | **ATTRIBUTE_SampleType** | **ATTRIBUTE_FractionDescription** | **ATTRIBUTE_Phenotype** | **ATTRIBUTE_Alias** |
| --- | --- | --- | --- | --- |
| 20220422_C18_95t05_l_Sample_00_MS2-r001.mzML | control | MeOH | NA | Solvent (MeOH) |
| 20220422_C18_95t05_l_Sample_00_MS2-r002.mzML | control | MeOH | NA | Solvent (MeOH) |
| 20220422_C18_95t05_l_Sample_00_MS2-r003.mzML | control | MeOH | NA | Solvent (MeOH) |
| 20220422_C18_95t05_l_Sample_02_MS2-r001.mzML | control | DCM_MeOH | NA | Solvent (MeOH:DCM) |
| 20220422_C18_95t05_l_Sample_02_MS2-r002.mzML | control | DCM_MeOH | NA | Solvent (MeOH:DCM) |
| 20220422_C18_95t05_l_Sample_02_MS2-r003.mzML | control | DCM_MeOH | NA | Solvent (MeOH:DCM) |
| 20220422_C18_95t05_l_Sample_03_MS2-r001.mzML | fraction | hexanes | rods | F1 |
| 20220422_C18_95t05_l_Sample_03_MS2-r002.mzML | fraction | hexanes | rods | F1 |
| 20220422_C18_95t05_l_Sample_03_MS2-r003.mzML | fraction | hexanes | rods | F1 |
| 20220422_C18_95t05_l_Sample_04_MS2-r001.mzML | fraction | hexanes_EtOAc | mixed | F2 |
| 20220422_C18_95t05_l_Sample_04_MS2-r002.mzML | fraction | hexanes_EtOAc | mixed | F2 |
| 20220422_C18_95t05_l_Sample_04_MS2-r003.mzML | fraction | hexanes_EtOAc | mixed | F2 |
| 20220422_C18_95t05_l_Sample_05_MS2-r001.mzML | crude_extract | EtOAc Extract | disks | Extract |
| 20220422_C18_95t05_l_Sample_05_MS2-r002.mzML | crude_extract | EtOAc Extract | disks | Extract |
| 20220422_C18_95t05_l_Sample_05_MS2-r003.mzML | crude_extract | EtOAc Extract | disks | Extract |
| 20220422_C18_95t05_l_Sample_06_MS2-r001.mzML | fraction | DCM_MeOH_pt1 | mixed | F5 |
| 20220422_C18_95t05_l_Sample_06_MS2-r002.mzML | fraction | DCM_MeOH_pt1 | mixed | F5 |
| 20220422_C18_95t05_l_Sample_06_MS2-r003.mzML | fraction | DCM_MeOH_pt1 | mixed | F5 |
| 20220422_C18_95t05_l_Sample_07_MS2-r001.mzML | fraction | DCM_MeOH_pt2 | disks | F7 |
| 20220422_C18_95t05_l_Sample_07_MS2-r002.mzML | fraction | DCM_MeOH_pt2 | disks | F7 |
| 20220422_C18_95t05_l_Sample_07_MS2-r003.mzML | fraction | DCM_MeOH_pt2 | disks | F7 |
| 20220422_C18_95t05_l_Sample_08_MS2-r001.mzML | fraction | EtOAc | disks | F3 |
| 20220422_C18_95t05_l_Sample_08_MS2-r002.mzML | fraction | EtOAc | disks | F3 |
| 20220422_C18_95t05_l_Sample_08_MS2-r003.mzML | fraction | EtOAc | disks | F3 |
| 20220422_C18_95t05_l_Sample_09_MS2-r001.mzML | fraction | DCM | rods | F4 |
| 20220422_C18_95t05_l_Sample_09_MS2-r002.mzML | fraction | DCM | rods | F4 |
| 20220422_C18_95t05_l_Sample_09_MS2-r003.mzML | fraction | DCM | rods | F4 |
| 20220422_C18_95t05_l_Sample_10_MS2-r001.mzML | fraction | YellowBand | disks | F6 |
| 20220422_C18_95t05_l_Sample_10_MS2-r002.mzML | fraction | YellowBand | disks | F6 |
| 20220422_C18_95t05_l_Sample_10_MS2-r003.mzML | fraction | YellowBand | disks | F6 |

**Detailed schematics for synthesis of DKP molecules**

*General*

Unless otherwise noted: all reactions were conducted using oven (135 ºC) dried glassware under a dry nitrogen atmosphere using standard Schlenk-line techniques unless otherwise noted. Methanol and dichloromethane used in reactions were purified, degassed, and dispensed under argon using a Solvent Purification System (Pure Process Technology). All work-up and purification procedures were conducted using reagent grade solvents (purchase from Fisher, VWR, or Sigma-Aldrich) in air. Triethylamine (Et_3_N) and diisopropylethyl amine (DIPEA) were used without additional purification. Silicycle SiliaFlash P60 (230-400 mesh) was used for purifications via standard column chromatography techniques. Microwave reactions were carried out in a Biotage Initiator+ microwave reactor in 2.0-5.0 mL Biotage-brand microwave vials sealed via crimping with a metal cap. ^1^H and ^13^C NMR spectra were recorded on a Bruker 400. Chemical shifts (δ) are reported in parts per million (ppm) relative to internal residual solvent peaks from indicated deuterated solvents. Coupling constants (J) are reported in Hertz (Hz) and are rounded to the nearest 0.1 Hz. Multiplicities are defined as: s = singlet, d = doublet, t = triplet, q = quartet, m = multiplet, dd = doublet of doublets, dt = doublet of triplets, ddd = doublet of doublet of doublets, dddd = doublet of doublet of doublet of doublets, br = broad, app = apparent, par = partial.

*General Scheme for Synthesis of Diketopiperazines* ***6***

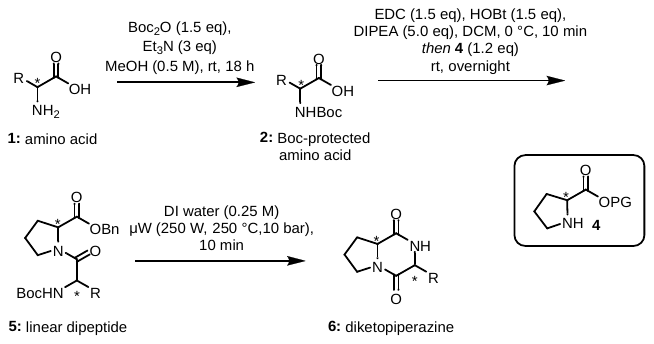

**Synthesis of Boc-Protected Amino Acids**

*General Procedure for the Preparation of Boc-Protected Amino Acids* ***2a-f*** ^75^

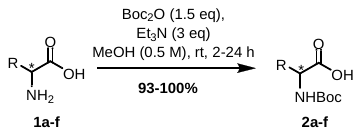

A 20 mL scintillation vial equipped with a stir bar was charged with starting amino acid **1a-f** (1.0 eq) and the appropriate quantity of methanol to give a 0.5 M solution. Triethylamine (3.0 eq) was then added, followed by Boc_2_O (1.5 eq). The reaction mixture was stirred until starting material had been consumed, monitored by TLC. Upon reaction completion, volatiles were removed by rotary evaporation, and the product was carried forward crude with triethylamine to the next step. Yields were determined with 4-fluorophenol as NMR internal standard.

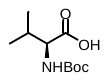
**2a,** *N*-Boc-L-Val**:** clear syrup, 99%. R_f_ = 0.86 (5:1 reagent alcohol:water); ^1^H NMR (CDCl_3_, 400 MHz) δ 10.26 (s, 1H), 5.33 (d, *J* = 8.4 Hz, 1H), 4.05 (dd, *J* = 8.5, 4.0 Hz, 1H), *3.02 (q, 5H, Et_3_N),* 2.24 – 2.12 (m, 1H), 1.42 (s, 9H), *1.24 (t, 8H, Et_3_N),* 0.95 (d, *J* = 6.9 Hz, 3H), 0.87 (d, *J* = 6.8 Hz, 3H).

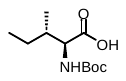
**2b,** *N*-Boc-L-Ile**:** clear syrup, 99%. R_f_ = 0.85 (5:1 reagent alcohol:water); ^1^H NMR (CDCl_3_, 400 MHz) δ 9.03 (s, 1H), 5.25 (d, J = 8.8 Hz, 1H), 4.18 (dd, J = 8.8, 4.6 Hz, 1H), *3.08 (q, 2H, Et_3_N),* 1.93 – 1.77 (m, 1H), 1.52 – 1.45 (m, 1H), 1.45 – 1.41 (m, 9H), *1.25 (t, 3H, Et_3_N),* 1.22 – 1.11 (m, 1H), 0.96 – 0.88 (m, 6H).

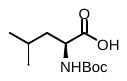
**2c,** *N*-Boc-L-Leu**:** clear syrup, quant. R_f_ = 0.77 (5:1 reagent alcohol:water); ^1^H NMR (CDCl_3_, 400 MHz) δ 8.54 (s, 1H), 5.16 (d, J = 9.1 Hz, 1H), 4.21 (td, J = 8.8, 4.7 Hz, 1H), *3.08 (q, 3H, Et_3_N),*1.81 – 1.70 (m, 1H), 1.70 – 1.59 (m, 1H), 1.52 – 1.47 (m, 1H), *1.25 (t, 4H, Et_3_N),* 1.43 (s, 9H), 0.94 (t, J = 7.2 Hz, 6H).

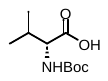
 **2d,** *N*-Boc-D-Val: clear syrup, 97%. R_f_ = 0.78 (5:1 reagent alcohol:water); ^1^H NMR (CDCl_3_, 400 MHz): δ 9.93 (s, 1H), 5.33 (d, J = 8.4 Hz, 1H), 4.05 (dd, J = 8.6, 4.0 Hz, 1H), *3.02 (q, 5H, Et_3_N),* 2.23 – 2.04 (m, 1H), 1.42 (s, 9H), *1.24 (t, 8H, Et_3_N),* 0.95 (d, J = 6.9 Hz, 3H), 0.87 (d, J = 6.8 Hz, 3H).

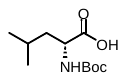
**2e,** *N*-Boc-D-Leu: clear syrup, 93%. R_f_ = 0.86 (5:1 reagent alcohol:water); ^1^H NMR (CDCl_3_, 400 MHz): δ 9.90 (s, 1H), 5.24 (d, J = 8.2 Hz, 1H), 4.23 – 4.09 (m, 1H), *3.03 (q, 4H, Et_3_N)*, 1.80 – 1.61 (m, 2H), 1.51 – 1.47 (m, 1H), 1.42 (s, 9H), *1.24 (t, 8H, Et_3_N)*, 0.95 (d, J = 6.3 Hz, 3H), 0.92 (d, J = 6.5 Hz, 3H).

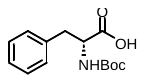
**2f**, *N*-Boc-D-Phe: clear yellowish syrup, 95%. R_f_ = 0.87 (5:1 reagent alcohol:water); ^1^H NMR (CDCl_3_, 400 MHz) δ 7.22 – 7.13 (m, 5H), 5.38 (d, J = 7.0 Hz, 1H), 4.35 (q, J = 5.7 Hz, 1H), 3.24 (dd, J = 13.6, 5.3 Hz, 1H), 3.09 (dd, J = 13.5, 5.3 Hz, 1H), 2.99 – 2.87 (*m, 6H,* *Et_3_N)*, 1.40 (s, 9H), 1.20 (*t, J = 7.3 Hz, 10H,* *Et_3_N)*.

*Procedure for the Synthesis of* N*-Boc-D-Tyr (****2g****)*^76^

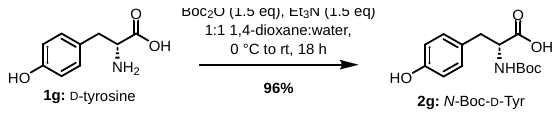

To a 50 mL teardrop-shaped round bottom flask containing D-tyrosine (1.0 g, 5.52 mmol, 1.0 eq) was added a prepared solution of 1:1 dioxane:water (18.08 mL) and a small stir bar. While stirring, triethylamine (1.15 mL, 8.28 mmol, 1.5 eq) was added. The solution was then cooled to 0 °C, and a solution of Boc_2_O (1.81 g in 4 mL 1:1 dioxane:water, 8.28 mmol, 1.5 eq) was added in one portion. The reaction mixture was stirred for 30 minutes, at which point the ice bath was removed and the reaction was allowed to stir at room temperature overnight.

After the reaction appeared complete by TLC, volatiles were removed by rotary evaporation, and the residue was taken back up in 40 mL of 1:1 ethyl acetate:DI water. The aqueous layer was extracted with ethyl acetate (3 x 20 mL), acidified to pH 1 with 1N HCl, and extracted again with ethyl acetate (3 x 20 mL). The combined organic layers were washed with brine (1 x 20 mL), dried over magnesium sulfate, and concentrated to give the product as a sticky white foam.

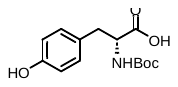
**2g,** *N*-Boc-D-Tyr: sticky white foam, 96%. R_f_ = 0.75 (5:1 reagent alcohol:water); ^1^H NMR (CDCl_3_, 400 MHz): δ 6.99 (d, J = 8.7 Hz, 2H), 6.72 (d, J = 8.3 Hz, 2H), 5.06 (d, J = 7.7 Hz, 1H), 4.54 (s, 1H), 3.08 – 2.98 (m, 2H), 1.42 (s, 9H).

**Synthesis of Protected Proline Starting Materials**

*Procedure for the Synthesis of L-Pro-OBn* ***4a***

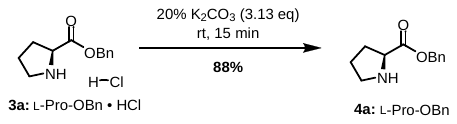

A 20 mL scintillation vial equipped with a stir bar was charged with L-proline benzyl ester hydrochloride **3** (1.0 eq) and an aqueous solution of potassium carbonate (3.13 eq, 20% w/w). The solution was stirred for 15 minutes or until all solids had dissolved. The solution was extracted with ethyl acetate five times; the combined organic layers were washed once with brine, dried over magnesium sulfate, and concentrated via rotary evaporation. Amine **4a** was obtained as a fluid yellow oil.

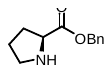
**4a,** L-Pro-OBn: fluid yellow oil, 88%. ^1^H NMR (CDCl_3_, 400 MHz) δ 7.43 – 7.25 (m, 6H), 5.18 (d, *J* = 1.2 Hz, 2H), 3.87 (dd, *J* = 8.6, 5.7 Hz, 1H), 3.19 – 3.07 (m, 2H), 2.97 (dt, *J* = 10.3, 6.7 Hz, 1H), 2.17 (dtd, *J* = 12.5, 8.3, 6.9 Hz, 1H), 1.90 (ddt, *J* = 12.4, 8.6, 6.0 Hz, 1H), 1.80 (ddq, *J* = 13.0, 8.2, 6.2 Hz, 2H).

*Procedure for the Synthesis of L-Pro-OMe Hydrochloride* ***3c***^77^

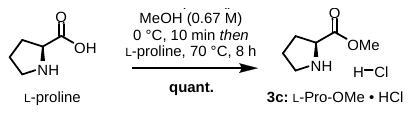

An oven-dried three-neck round bottom flask containing an oven-dried stir bar was fitted with an oven-dried reflux condenser. The reaction set-up was sealed with septa and allowed to cool to room temperature under nitrogen. To the flask was added anhydrous methanol (32.4 mL, 0.67 M). The flask was cooled in an ice bath before adding acetyl chloride (4.65 mL, 65.14 mmol, 3.0 eq). The mixture was allowed to stir on ice for 10 minutes. At this point, L-proline (2.50 g, 21.71 mmol, 1.0 eq) was added in one portion under positive pressure of nitrogen. The ice bath was removed and replaced with an oil bath, and the reaction mixture was refluxed at 70 °C for 8 hours, or until reaction completion was observed by TLC. Volatiles were removed via rotary evaporation, with the vacuum pump protected with a bubbler containing 10% KOH in water.

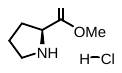
 **3c,** L-Pro-OMe • HCl: quant. yield R_f_ = 0.67 (5:1 reagent alcohol:water); ^1^H NMR (D_2_O, 400 MHz) δ 4.59 – 4.42 (m, 1H), 3.87 (s, 3H), 3.46 (qt, J = 11.5, 7.1 Hz, 2H), 2.47 (ddt, J = 15.5, 8.7, 4.4 Hz, 1H), 2.27 – 2.15 (m, 1H), 2.14 – 2.04 (m, 2H).

**Synthesis of Linear Dipeptides**

*General Procedures for the Synthesis of Linear Dipeptides* ***5a-f****, derived from L-Proline*^78,79^

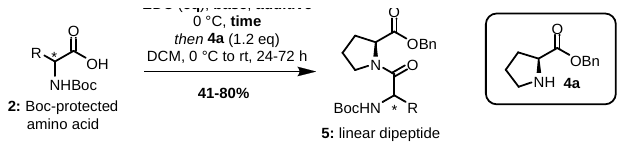

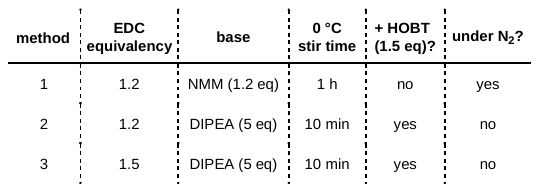

**Method 1:** To a 50 mL round bottom flask containing a stir bar was added EDC (1.2 eq) and *N*-methylmorpholine (1.2 eq). The contents were placed under nitrogen atmosphere (vacuumed and backfilled x 3). Dry DCM was added, and the contents were stirred to dissolution. A 1.0 M solution of the appropriate Boc-protected amino acid (**2a-f**) in dry DCM (1.0 eq) was added slowly, and the solution was stirred at 0 °C for 1 hour. At this point, a 1.0 M solution of L-Pro-OBn **4a** (1.2 eq) in dry DCM was added to the vial, and the solution was stirred at 0 °C for 3 hours and at room temperature until the limiting reagent appeared consumed by TLC. The total amount of DCM used in the reaction was the appropriate quantity to make the Boc-protected amino acid 0.11 M in the final solution

Upon reaction completion as determined by TLC, the reaction mixture was diluted with DCM and extracted three times with 10% aqueous citric acid. The organic layer was dried over magnesium sulfate and concentrated by rotary evaporation. The crude product was purified by column chromatography with 3:1 hexanes:ethyl acetate as eluent.

**Method 2:** To a 50 mL round bottom flask containing a stir bar was added the relevant Boc-protected amino acid (**2a-f,** 1.0 eq), a stir bar, and wet DCM. The acid was stirred on ice to dissolution. EDC (1.2 eq), HOBt (1.5 eq), and DIPEA (5 eq) were added to the solution, which was stirred on ice for 10 minutes. At this time, the amine **4a** (1.2 eq) was added as a solution in the appropriate amount of DCM so that the Boc-protected amino acid was 0.2 M in the final solution. The ice bath was removed and the reaction was allowed to stir at room temperature at least overnight.

Upon reaction completion as determined by TLC, volatiles were removed by rotary evaporation, and the crude residue was taken back up in ethyl acetate. This solution was extracted with 10% aqueous citric acid three times, saturated sodium bicarbonate three times, and saturated brine once. The organic layer was dried over magnesium sulfate, filtered, and concentrated by rotary evaporation. The crude product was purified by column chromatography with 3:1 hexanes:ethyl acetate as eluent.

**Method 3:** To a 50 mL round bottom flask containing a stir bar was added the relevant Boc-protected amino acid (**2a-f,** 1.0 eq), a stir bar, and wet DCM. The acid was stirred on ice to dissolution. EDC (1.5 eq), HOBt (1.5 eq), and DIPEA (5 eq) were added to the solution, which was stirred on ice for 10 minutes. At this time, the amine **4a** (1.2 eq) was added as a solution in the appropriate amount of DCM so that the Boc-protected amino acid was 0.2 M in the final solution. The ice bath was removed and the reaction was allowed to stir at room temperature at least overnight.

Upon reaction completion as determined by TLC, volatiles were removed by rotary evaporation, and the crude residue was taken back up in ethyl acetate. This solution was extracted with 10% aqueous citric acid three times, saturated sodium bicarbonate three times, and saturated brine once. The organic layer was dried over magnesium sulfate, filtered, and concentrated by rotary evaporation. The crude product was purified by column chromatography with 3:1 hexanes:ethyl acetate as eluent.

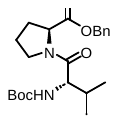
**5a,** *N*-Boc-L-Val-L-Pro-OBn: Synthesized via **Method 3**. clear syrup, 80%. R_f_ = 0.35, (2:1 hexanes:ethyl acetate); ^1^H NMR (CDCl_3_, 400 MHz): δ 7.37 – 7.27 (m, 5H), 5.36 (d, J = 9.4 Hz, 1H), 5.16 (dd, J = 12.4, 5.1 Hz, 2H), 4.56 (dd, J = 8.6, 4.6 Hz, 1H), 4.26 (dd, J = 9.5, 6.5 Hz, 1H), 3.78 (dt, J = 9.6, 6.4 Hz, 1H), 3.63 (qd, J = 7.1, 4.4 Hz, 1H), 2.26 – 2.09 (m, 1H), 2.07 – 1.89 (m, 4H), 1.41 (s, 9H), 0.97 (d, J = 6.8 Hz, 3H), 0.89 (d, J = 6.7 Hz, 3H).

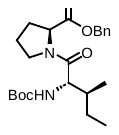
**5b,** *N-*Boc-L-Ile-L-Pro-OBn: Synthesized via **Method 2.** clear syrup, 41%. R_f_ = 0.30 (3:1 hexanes:ethyl acetate); ^1^H NMR (CDCl_3_, 400 MHz): δ 7.35 – 7.20 (m, 6H) (*overintegrates by 1*), 5.18 (d, J = 9.7 Hz, 1H), 5.12 (dd, J = 12.6, 2.5 Hz, 2H), 4.54 (dd, J = 8.5, 4.3 Hz, 1H), 4.26 (dd, J = 9.5, 7.3 Hz, 1H), 3.78 (dt, J = 9.5, 6.2 Hz, 1H), 3.62 (dtd, J = 9.9, 5.9, 2.6 Hz, 1H), 2.25 – 2.08 (m, 1H), 2.06 – 1.86 (m, 3H), 1.71 (dtt, J = 13.5, 9.8, 4.9 Hz, 1H), 1.55 (dqd, J = 14.9, 7.4, 3.1 Hz, 1H), 1.39 (s, 9H), 1.08 (ddt, J = 16.6, 14.1, 7.2 Hz, 1H), 0.93 (d, J = 6.8 Hz, 3H), 0.84 (t, J = 7.5 Hz, 3H).

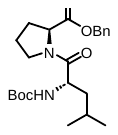
**5c,** *N-*Boc-L-Leu-L-Pro-OBn: Synthesized via **Method 1.** pale yellow syrup, 60%. R_f_ = 0.24 (3:1 hexanes:ethyl acetate), 0.91 (5:1 reagent alcohol:water); ^1^H NMR (CDCl_3_, 400 MHz): δ 7.38 – 7.30 (m, 5H), 5.20 (d, J = 9.2 Hz, 1H), 5.19 (d, J = 12.3 Hz, 1H), 5.09 (d, J = 12.3 Hz, 1H), 4.59 (dd, J = 8.7, 4.0 Hz, 1H), 4.46 (td, J = 8.8, 5.6 Hz, 1H), 3.77 (dt, J = 9.7, 6.6 Hz, 1H), 3.58 (dt, J = 9.5, 6.4 Hz, 1H), 2.28 – 2.15 (m, 1H), 2.08 – 1.93 (m, 3H), 1.73 (septet, J = 6.8 Hz, 1H), 1.48 – 1.43 (m, 2H), 1.42 (s, 9H), 0.96 (d, J = 6.5 Hz, 3H), 0.90 (d, J = 6.8 Hz, 3H).

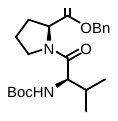
**5d,** *N*-Boc-D-Val-L-Pro-OBn: Synthesized via **Method 2.** off-white solid, 64%. R_f_ = 0.93 (5:1 reagent alcohol:water), 0.33 (3:1 hexanes:ethyl acetate) ^1^H NMR (CDCl_3_, 400 MHz): δ 7.40 – 7.27 (m, 5H), 5.25 – 4.96 (m, 3H), 4.48 (dd, J = 8.6, 3.3 Hz, 1H), 4.33 (dd, J = 9.3, 6.2 Hz, 1H), 3.87 (td, J = 8.7, 7.5, 4.3 Hz, 1H), 3.63 – 3.50 (m, 1H), 2.24 – 1.78 (m, 5H), 1.50 – 1.39 (m, 9H), 1.00 – 0.86 (m, 6H).

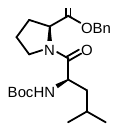
**5e,** *N-*Boc-D-Leu-L-Pro-OBn: Synthesized via **Method 2.** yellow-tinted syrup, 45%. R_f_ = 0.30 (3:1 hexanes:ethyl acetate), 0.88 (5:1 reagent alcohol:water); ^1^H NMR (CDCl_3_, 400 MHz): δ 7.40 – 7.27 (m, 6H) *(overintegrates by 1)*, 5.23 – 5.09 (m, 2H), 5.04 – 4.94 (m, 0.38H), 4.70 (s, 0.61H) *(impurity)*, 4.55 (dt, J = 9.3, 4.7 Hz, 1H), 4.47 (dd, J = 8.5, 3.4 Hz, 1H), 4.25 (td, J = 10.3, 3.6 Hz, 0.19H), 3.86 (ddd, J = 10.1, 7.5, 4.3 Hz, 1H), 3.55 (ddt, J = 30.5, 9.7, 6.9 Hz, 1H), 2.32 – 2.05 (m, 2H), 2.04 – 1.81 (m, 2H), 1.76 – 1.59 (m, 2H), 1.56 – 1.33 (m, 10H), 1.04 – 0.75 (m, 6H).

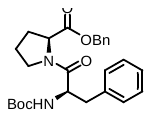
 **5f,** *N-*Boc-D-Phe-L-Pro-OBn: clear syrup, 50%. R_f_ = 0.186, (3:1 hexanes:ethyl acetate); ^1^H NMR (CDCl_3_, 400 MHz): δ 7.38 – 7.30 (m, 7H), 7.26 – 7.17 (m, 5H), 5.37 (d, J = 8.6 Hz, 1H), 5.23 – 5.06 (m, 2H), 4.64 (td, J = 9.1, 5.5 Hz, 1H), 4.35 (dd, J = 8.2, 4.0 Hz, 1H), 3.51 (ddd, J = 9.7, 7.3, 5.1 Hz, 1H), 3.06 (dd, J = 12.8, 5.5 Hz, 1H), 2.93 (dd, J = 12.8, 9.4 Hz, 1H), 2.62 (dt, J = 9.7, 6.9 Hz, 1H), 1.99 – 1.73 (m, 3H), 1.54 – 1.45 (m, 1H), 1.43 (s, 9H).

*Procedure for the Synthesis of Linear Dipeptide* ***5g*** ^80^

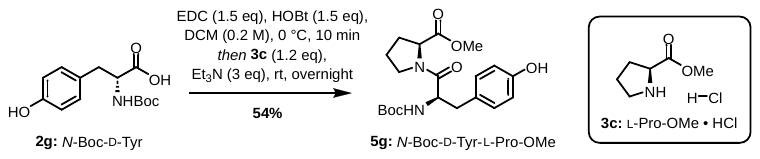

To a 50 mL round bottom flask containing a stir bar was added **2g** (843.9 mg, 3.00 mmol, 1.0 eq), a stir bar, and wet DCM (12 mL). The acid was stirred on ice to dissolution. EDC (698.6 mg, 4.50 mmol, 1.2 eq) and HOBt (608.1 mg, 4.50 mmol, 1.5 eq) were added to the solution, which was stirred on ice for 10 minutes. At this time, the amine **3c** (L-Pro-OMe • HCl) (596.2 mg, 3.60 mmol, 1.2 eq) was added as a solution in triethylamine (1.25 mL, 9.00 mmol, 1.2 eq) and DCM (3 mL). The ice bath was removed and the reaction was allowed to stir at room temperature at least overnight.

Upon reaction completion as determined by TLC, volatiles were removed by rotary evaporation, and the crude residue was taken back up in ethyl acetate (50 mL). This solution was extracted with 10% aqueous citric acid (3 x 50 mL), saturated sodium bicarbonate (3 x 50 mL), and saturated brine (1 x 50 mL). The organic layer was dried over magnesium sulfate, filtered, and concentrated by rotary evaporation. The crude product was purified by column chromatography with 1:1 hexanes:ethyl acetate as eluent.

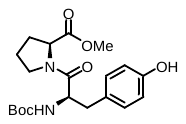
 **5g,** *N-*Boc-D-Tyr-L-Pro-OMe: white foam, 54%. R_f_ = 0.23 (1:1 hexanes:ethyl acetate); ^1^H NMR (CDCl_3_, 400 MHz): δ 7.06 – 7.02 (m, 2H), 6.75 – 6.69 (m, 2H), 5.55 (s, 1H), 5.37 (d, J = 8.7 Hz, 1H), 4.59 (td, J = 9.1, 5.4 Hz, 1H), 4.30 (dd, J = 8.5, 3.8 Hz, 1H), 3.71 (s, 3H), 3.56 (dt, J = 13.1, 6.5 Hz, 1H), 2.96 (dd, J = 13.1, 5.3 Hz, 1H), 2.85 (dd, J = 13.1, 9.3 Hz, 1H), 2.74 (dt, J = 9.8, 7.1 Hz, 1H), 1.96 (td, J = 9.3, 7.6, 4.1 Hz, 1H), 1.92 – 1.81 (m, 2H), 1.61 – 1.54 (m, 1H), 1.43 (s, 9H).

*General Procedures for the Synthesis of Linear Dipeptides* ***5h-p****, derived from D-Proline ^5,6^*

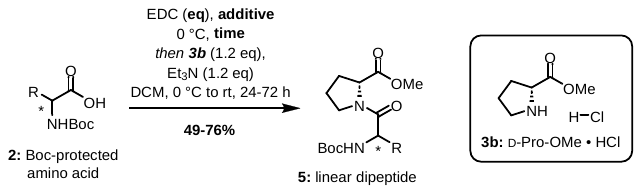

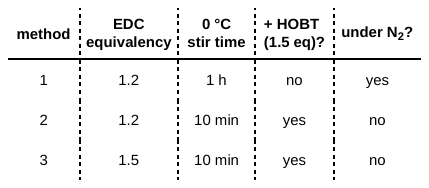

**Method 1:** To a 20 mL scintillation vial was added D-Pro-OMe • HCl (1.2 eq) and a small stir bar, which were then placed under nitrogen atmosphere (vacuumed and backfilled x 3). To this vial was added dry DCM and dry triethylamine (1.2 eq). The contents of the vial were stirred at room temperature for at least one hour until all solids were dissolved.

To a separate 50 mL round bottom flask containing a stir bar was added EDC (1.2 eq). The contents were placed under nitrogen atmosphere (vacuumed and backfilled x 3). Dry DCM was added, and the contents were stirred to dissolution. A 1.0 M solution of the appropriate Boc-protected amino acid (**2a-f**, 1.0 eq) in dry DCM was added slowly, and the solution was stirred at 0 °C for 1 hour. At this point, the contents of the first vial were added to the flask, and the solution was stirred at 0 °C for 3 hours and at room temperature until the limiting reagent appeared consumed by TLC. The total amount of DCM used in the reaction was the appropriate quantity to make the Boc-protected amino acid 0.11 M in the final solution.

Upon reaction completion as determined by TLC, the reaction mixture was diluted with DCM and extracted three times with 10% aqueous citric acid. The organic layer was dried over magnesium sulfate and concentrated by rotary evaporation. The crude product was purified by column chromatography with 3:1 hexanes:ethyl acetate as eluent.

**Method 2:** To a 50 mL round bottom flask containing a stir bar was added the relevant Boc-protected amino acid (**2a-f,** 1.0 eq), a stir bar, and wet DCM. The acid was stirred on ice to dissolution. EDC (1.2 eq) and HOBt (1.5 eq) were added to the solution, which was stirred on ice for 10 minutes. At this time, the amine D-Pro-OMe • HCl (1.2 eq) was added as a solution in triethylamine (1.2 eq) and the appropriate amount of DCM so that the Boc-protected amino acid was 0.2 M in the final solution. The ice bath was removed and the reaction was allowed to stir at room temperature at least overnight.

Upon reaction completion as determined by TLC, volatiles were removed by rotary evaporation, and the crude residue was taken back up in ethyl acetate. This solution was extracted with 10% aqueous citric acid three times, saturated sodium bicarbonate three times, and saturated brine once. The organic layer was dried over magnesium sulfate, filtered, and concentrated by rotary evaporation. The crude product was purified by column chromatography with 3:1 hexanes:ethyl acetate as eluent.

**Method 3:** To a 50 mL round bottom flask containing a stir bar was added the relevant Boc-protected amino acid (**2a-f,** 1.0 eq), a stir bar, and wet DCM. The acid was stirred on ice to dissolution. EDC (1.5 eq) and HOBt (1.5 eq) were added to the solution, which was stirred on ice for 10 minutes. At this time, the amine D-Pro-OMe • HCl (1.2 eq) was added as a solution in triethylamine (1.2 eq) and the appropriate amount of DCM so that the Boc-protected amino acid was 0.2 M in the final solution. The ice bath was removed and the reaction was allowed to stir at room temperature at least overnight.

Upon reaction completion as determined by TLC, volatiles were removed by rotary evaporation, and the crude residue was taken back up in ethyl acetate. This solution was extracted with 10% aqueous citric acid three times, saturated sodium bicarbonate three times, and saturated brine once. The organic layer was dried over magnesium sulfate, filtered, and concentrated by rotary evaporation. The crude product was purified by column chromatography with 3:1 hexanes:ethyl acetate as eluent.

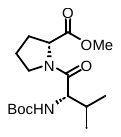
**5h,** N-Boc-L-Val-D-Pro-OMe: Synthesized via **Method 1.** yellow-tinted syrup, 54%. R_f_ = 0.18 (3:1 hexanes:ethyl acetate), 0.95 (5:1 reagent alcohol:water); ^1^H NMR (CDCl_3_, 400 MHz): δ 5.23 (d, J = 9.3 Hz, 1H), 4.42 (dd, J = 8.4, 3.5 Hz, 1H), 4.32 (dd, J = 9.4, 6.3 Hz, 1H), 3.88 (ddd, J = 9.7, 5.0, 2.9 Hz, 1H), 3.73 (d, J = 10.8 Hz, 3H), 3.62 – 3.51 (m, 1H), 2.25 – 2.05 (m, 2H), 2.05 – 1.89 (m, 3H), 1.51 – 1.36 (m, 9H), 1.01 – 0.84 (m, 6H).

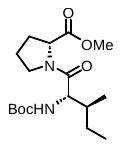
**5i,** *N*-Boc-L-Ile-D-Pro-OMe: Synthesized via **Method 1.** clear, partially crystalline syrup, 53%. R_f_ = 0.24 (3:1 hexanes:ethyl acetate), 0.95 (5:1 reagent alcohol:water); ^1^H NMR (CDCl_3_, 400 MHz): δ 5.16 (d, J = 9.5 Hz, 1H), 4.42 (dd, J = 8.3, 3.6 Hz, 1H), 4.33 (dd, J = 9.5, 7.0 Hz, 1H), 3.91 (ddd, J = 9.5, 7.1, 4.4 Hz, 1H), 3.73 (d, J = 10.0 Hz, 3H), 3.57 (ddt, J = 14.7, 9.9, 4.5 Hz, 1H), 2.27 – 2.04 (m, 2H), 1.99 (dddd, J = 21.0, 10.5, 4.1, 2.6 Hz, 2H), 1.70 (ddt, J = 13.4, 6.9, 3.3 Hz, 1H), 1.56 (ddd, J = 13.5, 7.5, 3.4 Hz, 1H), 1.42 (d, J = 7.2 Hz, 9H), 1.11 (ddt, J = 14.1, 9.3, 7.2 Hz, 1H), 0.95 – 0.81 (m, 6H).

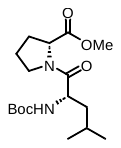
**5j,** *N*-Boc-L-Leu-D-Pro-OMe: Synthesized via **Method 1.** colorless syrup, 52%. R_f_ = 0.22 (3:1 hexanes:ethyl acetate), 0.89 (5:1 reagent alcohol:water); ^1^H NMR (CDCl_3_, 400 MHz): δ 5.20 (d, J = 9.1 Hz, 1H), 4.52 (td, J = 9.3, 4.4 Hz, 1H), 4.41 (dd, J = 8.4, 3.7 Hz, 1H), 3.87 (ddd, J = 9.9, 7.3, 4.6 Hz, 1H), 3.73 (d, J = 10.0 Hz, 3H), 3.62 – 3.46 (m, 1H), 2.32 – 2.06 (m, 2H), 2.03 – 1.86 (m, 2H), 1.77 – 1.62 (m, 1H), 1.61 – 1.45 (m, 2H), 1.42 (d, J = 8.3 Hz, 9H), 1.01 – 0.84 (m, 6H).

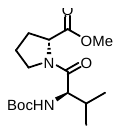
**5k,** *N*-Boc-D-Val-D-Pro-OMe): Synthesized via **Method 3.** clear syrup, 74%. R_f_ = 0.37 (3:1 hexanes:ethyl acetate), 0.61 (5:1 reagent alcohol:water); ^1^H NMR (CDCl_3_, 400 MHz) δ 5.19 (d, *J* = 9.4 Hz, 1H), 4.52 (dd, *J* = 8.7, 4.8 Hz, 1H), 4.28 (dd, *J* = 9.4, 6.3 Hz, 1H), 3.84 – 3.74 (m, 1H), 3.71 (s, 3H), 3.70 – 3.60 (m, 1H), 2.29 – 2.16 (m, 1H), 2.11 – 1.85 (m, 4H), 1.42 (s, 9H), 1.03 (d, *J* = 6.8 Hz, 3H), 0.94 (d, *J* = 6.8 Hz, 3H).

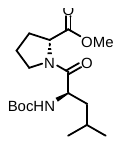
**5l,** *N*-Boc-D-Leu-D-Pro-OMe: Synthesized via **Method 3.** clear syrup, 70%. R_f_ = 0.23 (3:1 hexanes:ethyl acetate), 0.71 (5:1 reagent alcohol:water); ^1^H NMR (CDCl_3_, 400 MHz): δ 5.10 (d, *J* = 9.3 Hz, 1H), 4.53 (dd, *J* = 8.4, 4.4 Hz, 1H), 4.47 (td, *J* = 8.7, 5.6 Hz, 1H), 3.82 – 3.72 (m, 1H), 3.72 (s, 3H), 3.60 (ddd, *J* = 9.5, 7.3, 5.6 Hz, 1H), 2.29 – 2.15 (m, 1H), 2.12 – 1.90 (m, 3H), 1.77 (dp, *J* = 13.4, 6.7 Hz, 1H), 1.49 (td, *J* = 6.4, 5.9, 3.4 Hz, 2H), 1.42 (s, 9H), 1.00 (d, *J* = 6.5 Hz, 3H), 0.96 (d, *J* = 6.7 Hz, 3H).

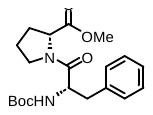
**5m,** *N*-Boc-L-Phe-D-Pro-OMe: Synthesized via **Method 3** using commercially available *N*-Boc-L-Phe as starting material**.** white foam, 76%. R_f_ = 0.16 (3:1 hexanes:ethyl acetate), 0.79 (5:1 reagent alcohol:water); ^1^H NMR (CDCl_3_, 400 MHz): δ 7.28 (dt, J = 4.5, 2.8 Hz, 1H), 7.25 – 7.13 (m, 4H), 5.36 (d, J = 8.8 Hz, 1H), 4.63 (td, J = 9.1, 5.5 Hz, 1H), 4.29 (dd, J = 8.0, 4.0 Hz, 1H), 3.71 (s, 3H), 3.52 (ddd, J = 9.9, 7.4, 5.3 Hz, 1H), 3.05 (dd, J = 13.0, 5.5 Hz, 1H), 2.92 (dd, J = 12.8, 9.4 Hz, 1H), 2.63 (dt, J = 9.5, 6.7 Hz, 1H), 1.98 – 1.76 (m, 3H), 1.51 (td, J = 7.2, 5.3 Hz, 1H), 1.43 (s, 9H).

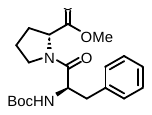
**5n,** *N*-Boc-D-Phe-D-Pro-OMe: Synthesized via **Method 3.** yellowish syrup, 50%. R_f_ = 0.15 (3:1 hexanes:ethyl acetate), 0.68 (5:1 reagent alcohol:water); ^1^H NMR (CDCl_3_, 400 MHz): δ 7.34 – 7.27 (m, 3H), 7.26 – 7.19 (m, 2H), 5.24 (d, J = 8.7 Hz, 1H), 4.64 (dt, J = 8.7, 6.8 Hz, 1H), 4.50 (dd, J = 8.5, 4.1 Hz, 1H), 3.74 (s, 2H), 3.69 (s, 1H), 3.67 – 3.49 (m, 1H), 3.18 (dt, J = 12.8, 6.5 Hz, 1H), 3.07 (td, J = 14.7, 14.2, 6.0 Hz, 1H), 2.96 – 2.84 (m, 1H), 2.16 (ttd, J = 11.5, 7.8, 6.8, 4.1 Hz, 1H), 1.93 (dq, J = 11.8, 6.7, 6.2 Hz, 3H), 1.42 (s, 2H), 1.38 (s, 7H).

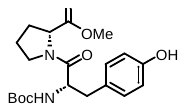
**5o,** *N*-Boc-L-Tyr-D-Pro-OMe: Synthesized via **Method 3** using commercially available *N*-Boc-L-Tyr as starting material**.** white foam, 66%. R_f_ = 0.04 (3:1 hexanes:ethyl acetate); ^1^H NMR (CDCl_3_, 400 MHz) δ 7.02 (dd, J = 18.3, 8.4 Hz, 2H), 6.72 (d, J = 8.4 Hz, 2H), 5.61 (s, 1H), 5.37 (d, J = 8.7 Hz, 1H), 4.59 (td, J = 8.9, 5.3 Hz, 1H), 4.30 (dd, J = 8.2, 3.8 Hz, 1H), 3.71 (s, 3H), 3.66 – 3.50 (m, 1H), 2.96 (dd, J = 13.1, 5.4 Hz, 1H), 2.85 (dd, J = 13.0, 9.2 Hz, 1H), 2.76 (dt, J = 9.7, 7.0 Hz, 1H), 2.01 – 1.79 (m, 3H), 1.61 – 1.53 (m, 1H), 1.43 (s, 9H).

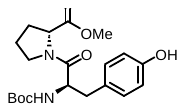
**5p,** *N*-Boc-D-Tyr-D-Pro-OMe: Synthesized via **Method 3.** white foam, 49%. R_f_ = 0.04 (3:1 hexanes:ethyl acetate); ^1^H NMR (CDCl_3_, 400 MHz): δ 7.15 – 7.00 (m, 2H), 6.75 – 6.61 (m, 2H), 5.94 (s, 1H), 5.25 (d, J = 8.9 Hz, 1H), 4.61 (dt, J = 9.0, 6.5 Hz, 1H), 4.51 (dd, J = 8.2, 3.9 Hz, 1H), 3.72 (d, J = 15.4 Hz, 3H), 3.61 (ddt, J = 17.3, 12.0, 6.6 Hz, 1H), 3.37 – 3.23 (m, 1H), 3.02 (dd, J = 14.0, 6.8 Hz, 1H), 2.84 (dt, J = 12.9, 7.4 Hz, 1H), 2.18 (td, J = 9.3, 8.5, 6.2 Hz, 1H), 1.95 (dp, J = 10.7, 6.3, 5.4 Hz, 3H), 1.41 (d, J = 12.9 Hz, 9H).

**Synthesis of Diketopiperazines** ***6a-p*** *^81^*

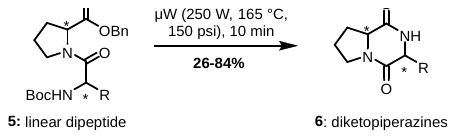

To a Biotage Initiator+ microwave reaction vial was added the relevant linear dipeptide, a small stir bar, and DI water (0.25 M). The tube was then capped, the cap was crimped, and the tube was placed in the Biotage Initiator+ reactor. The reaction was run for 10 minutes at 250 °C and 150 psi (10 bar) at 250 W.

Upon removal from the reactor, the reaction mixture in water was washed with DCM (3 x 10 mL), and the combined organic layer was dried over magnesium sulfate, filtered, and concentrated. The product was purified via silica plug with methanol/DCM as eluent.

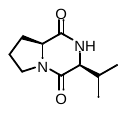
**6a,** cyclo-(L-Pro-L-Val): chunky white solid, 42%. ^1^H NMR (CDCl_3_, 400 MHz): δ 5.85 (s, 1H), 4.08 (ddd, J = 9.1, 6.7, 1.8 Hz, 1H), 3.93 (t, J = 2.2 Hz, 1H), 3.64 (dt, J = 12.1, 8.0 Hz, 1H), 3.54 (ddd, J = 11.9, 9.0, 3.2 Hz, 1H), 2.63 (heptd, J = 7.1, 2.7 Hz, 1H), 2.43 – 2.31 (m, 1H), 2.13 – 1.97 (m, 2H), 1.99 – 1.82 (m, 1H), 1.06 (d, J = 7.2 Hz, 3H), 0.91 (d, J = 6.8 Hz, 3H). ^13^C{^1^H} NMR (CDCl_3_, 101 MHz) δ 170.08, 165.02, 60.52, 58.97, 45.29, 28.68, 28.51, 22.51, 19.41, 16.20.

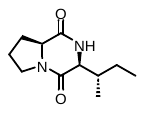
**6b,** cyclo-(L-Pro-L-Ile): white, yellow tinted solid, 46%. ^1^H NMR (CDCl_3_, 400 MHz): δ 5.89 (s, 1H), 4.07 (ddd, J = 9.4, 6.8, 1.9 Hz, 1H), 3.96 (t, J = 2.2 Hz, 1H), 3.71 – 3.48 (m, 2H), 2.44 – 2.24 (m, 2H), 2.12 – 1.97 (m, 2H), 1.97 – 1.82 (m, 1H), 1.43 (dqd, J = 13.3, 7.5, 3.9 Hz, 1H), 1.24 – 1.11 (m, 1H), 1.05 (d, J = 7.2 Hz, 3H), 0.93 (t, J = 7.4 Hz, 3H). ^13^C{^1^H} NMR (CDCl_3_, 101 MHz): δ 169.94, 165.10, 60.63, 58.93, 45.29, 35.44, 28.68, 24.19, 22.50, 16.08, 12.20.

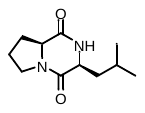
**6c,** cyclo-(L-Pro-L-Leu): white solid, 58%. ^1^H NMR (CDCl_3_, 400 MHz): δ 5.97 (s, 1H), 4.11 (ddd, J = 9.2, 7.0, 1.6 Hz, 1H), 4.01 (dt, J = 10.0, 2.6 Hz, 1H), 3.56 (dddd, J = 20.5, 12.0, 8.3, 3.8 Hz, 2H), 2.35 (dtd, J = 12.7, 6.9, 3.0 Hz, 1H), 2.20 – 1.97 (m, 3H), 1.96 – 1.84 (m, 1H), 1.81 – 1.68 (m, 1H), 1.52 (ddd, J = 14.5, 9.5, 5.0 Hz, 1H), 1.00 (d, J = 6.6 Hz, 3H), 0.95 (d, J = 6.6 Hz, 3H). ^13^C{^1^H} NMR (CDCl_3_, 101 MHz) δ 170.28, 166.30, 59.13, 53.54, 45.64, 38.77, 28.25, 24.86, 23.43, 22.89, 21.35.

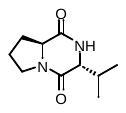
**6d,** cyclo-(L-Pro-D-Val): off-white solid, 56%. ^1^H NMR (CDCl_3_, 400 MHz): δ 6.49 – 6.31 (m, 1H), 4.09 (dd, J = 9.9, 6.5 Hz, 1H), 3.76 – 3.65 (m, 2H), 3.52 (ddd, J = 11.9, 8.9, 2.8 Hz, 1H), 2.41 (ddd, J = 12.2, 6.1, 4.4 Hz, 1H), 2.23 (pd, J = 6.9, 5.3 Hz, 1H), 2.03 (dddd, J = 15.6, 8.9, 3.3, 1.5 Hz, 1H), 1.98 – 1.79 (m, 2H), 1.04 (d, J = 6.9 Hz, 3H), 0.99 (d, J = 6.8 Hz, 3H). ^13^C{^1^H} NMR (101 MHz, Chloroform-*d*) δ 169.49, 165.36, 63.72, 58.45, 45.73, 33.28, 29.55, 22.08, 19.12, 17.67.

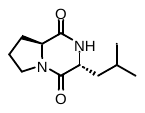
**6e,** cyclo-(L-Pro-D-Leu): white solid, 74%. ^1^H NMR (CDCl_3_, 400 MHz): δ 6.47 (s, 1H), 4.08 (dd, J = 9.5, 6.7 Hz, 1H), 3.92 (ddd, J = 9.6, 5.7, 4.3 Hz, 1H), 3.71 – 3.59 (m, 1H), 3.52 (ddd, J = 11.9, 8.7, 2.7 Hz, 1H), 2.46 – 2.34 (m, 1H), 2.11 – 1.96 (m, 2H), 1.96 – 1.83 (m, 1H), 1.83 – 1.72 (m, 1H), 1.70 – 1.59 (m, 2H), 0.99 (d, J = 6.5 Hz, 3H), 0.95 (d, J = 6.5 Hz, 3H). ^13^C{^1^H} NMR (101 MHz, Chloroform-d) δ 169.71, 166.53, 58.18, 56.46, 45.76, 42.73, 29.10, 24.60, 23.34, 23.16, 22.34, 21.53.

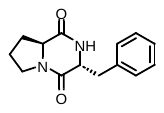
**6f,** cyclo-(L-Pro-D-Phe): white solid, 53%. ^1^H NMR (CD_3_OD 400 MHz): δ 7.38 – 7.28 (m, 3H), 7.25 – 7.16 (m, 2H), 4.22 (td, J = 4.8, 1.0 Hz, 1H), 3.62 – 3.50 (m, 1H), 3.39 – 3.34 (m, 1H), 3.21 (dd, J = 13.6, 4.8 Hz, 1H), 3.01 (dd, J = 13.7, 4.7 Hz, 1H), 2.64 (dd, J = 10.6, 6.2 Hz, 1H), 2.11 – 1.98 (m, 1H), 1.92 (dddd, J = 13.2, 8.6, 4.5, 1.9 Hz, 1H), 1.77 – 1.55 (m, 2H).

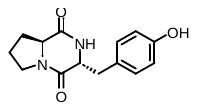
 **6g,** cyclo-(L-Pro-D-Tyr): yellow-white foamy solid, 73%. ^1^H NMR (CD_3_OD, 400 MHz): δ 7.07 – 6.94 (m, 2H), 6.76 – 6.67 (m, 2H), 4.14 (td, J = 4.5, 1.0 Hz, 1H), 3.54 (dt, J = 12.0, 8.4 Hz, 1H), 3.31 (p, J = 1.6 Hz, 7H) *(actually 1H - overlaps with MeOD peak)*, 3.11 (dd, J = 13.8, 4.3 Hz, 1H), 2.88 (dd, J = 13.9, 4.6 Hz, 1H), 2.61 (dd, J = 10.5, 6.3 Hz, 1H), 2.11 – 2.01 (m, 1H), 1.92 (pd, J = 7.4, 6.4, 4.1 Hz, 1H), 1.74 – 1.55 (m, 2H).

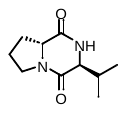
 **6h,** cyclo-(D-Pro-L-Val): white solid, 72%. ^1^H NMR (CDCl_3_, 400 MHz): δ 6.23 (s, 1H), 4.09 (dd, J = 9.9, 6.5 Hz, 1H), 3.77 – 3.65 (m, 2H), 3.52 (ddd, J = 11.9, 8.9, 2.8 Hz, 1H), 2.41 (ddd, J = 12.4, 6.3, 4.6 Hz, 1H), 2.24 (pd, J = 6.9, 5.3 Hz, 1H), 2.09 – 1.97 (m, 1H), 2.00 – 1.79 (m, 2H), 1.04 (d, J = 6.9 Hz, 3H), 0.99 (d, J = 6.8 Hz, 3H).  ^13^C{^1^H} NMR (101 MHz, Chloroform-d) δ 169.33, 165.32, 63.77, 58.45, 45.75, 33.27, 29.56, 22.08, 19.12, 17.65.

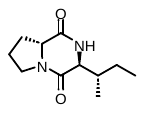
**6i,** cyclo-(D-Pro-L-Ile): off-white solid, 40%. ^1^H NMR (CDCl_3_, 400 MHz): δ 6.54 (s, 1H), 4.08 (dd, J = 9.9, 6.5 Hz, 1H), 3.78 (dd, J = 5.8, 4.0 Hz, 1H), 3.75 – 3.64 (m, 1H), 3.51 (ddd, J = 12.0, 8.9, 2.8 Hz, 1H), 2.40 (ddd, J = 13.0, 6.3, 4.7 Hz, 1H), 2.08 – 1.98 (m, 1H), 1.98 – 1.89 (m, 2H), 1.89 – 1.81 (m, 1H), 1.56 (dqd, J = 13.0, 7.4, 4.0 Hz, 1H), 1.31 – 1.16 (m, 1H), 1.01 (d, J = 6.9 Hz, 3H), 0.92 (t, J = 7.4 Hz, 3H). ^13^C{^1^H} NMR (101 MHz, Chloroform-d) δ 169.53, 165.36, 63.03, 58.50, 45.74, 39.79, 29.52, 24.66, 22.11, 15.41, 11.42.

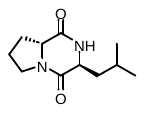
**6j,** cyclo-(D-Pro-L-Leu): white solid, 37%. ^1^H NMR (CDCl_3_, 400 MHz): δ 6.58 (d, *J* = 3.8 Hz, 1H), 4.07 (dd, *J* = 9.6, 6.8 Hz, 1H), 3.92 (ddd, *J* = 9.6, 5.7, 4.3 Hz, 1H), 3.71 – 3.59 (m, 1H), 3.51 (ddd, *J* = 12.0, 8.8, 2.6 Hz, 1H), 2.45 – 2.33 (m, 1H), 2.09 – 1.95 (m, 2H), 1.95 – 1.82 (m, 1H), 1.83 – 1.72 (m, 1H), 1.71 – 1.55 (m, 2H), 0.98 (d, *J* = 6.5 Hz, 3H), 0.95 (d, *J* = 6.5 Hz, 3H). ^13^C{^1^H} NMR (101 MHz, Chloroform-d) δ 169.51, 166.43, 58.18, 56.53, 45.78, 42.75, 29.12, 24.63, 23.17, 22.36, 21.54.

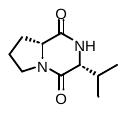
**6k,** cyclo-(D-Pro-D-Val): white solid, 84%. ^1^H NMR (400 MHz, Chloroform-*d*) δ 5.87 (s, 1H), 4.08 (ddd, *J* = 9.3, 6.7, 1.9 Hz, 1H), 3.93 (t, *J* = 2.5 Hz, 1H), 3.75 – 3.57 (m, 1H), 3.54 (ddd, *J* = 11.9, 8.9, 3.1 Hz, 1H), 2.63 (heptd, *J* = 7.0, 2.6 Hz, 1H), 2.43 – 2.31 (m, 1H), 2.13 – 1.97 (m, 2H), 1.99 – 1.80 (m, 1H), 1.06 (d, *J* = 7.3 Hz, 3H), 0.91 (d, *J* = 6.8 Hz, 3H). ^13^C{^1^H} NMR (101 MHz, Chloroform-*d*) δ 170.08, 165.02, 60.52, 58.97, 45.29, 28.68, 28.52, 22.51, 19.41, 16.20.

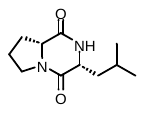
**6l,** cyclo-(D-Pro-D-Leu): white solid, 73%. ^1^H NMR (CDCl_3_, 400 MHz): δ 5.89 (s, 1H), 4.11 (ddd, J = 9.0, 7.0, 1.5 Hz, 1H), 4.05 – 3.97 (m, 1H), 3.66 – 3.47 (m, 2H), 2.35 (dtd, J = 12.7, 6.9, 3.0 Hz, 1H), 2.20 – 1.97 (m, 3H), 1.90 (ddtd, J = 12.8, 10.8, 8.7, 6.8 Hz, 1H), 1.74 (dddd, J = 16.4, 13.2, 6.7, 5.2 Hz, 1H), 1.52 (ddd, J = 14.5, 9.5, 5.0 Hz, 1H), 1.00 (d, J = 6.5 Hz, 3H), 0.95 (d, J = 6.6 Hz, 3H). ^13^C{^1^H} NMR (101 MHz, Chloroform-d) δ 170.25, 166.30, 59.14, 53.53, 45.65, 38.78, 28.26, 24.87, 23.43, 22.89, 21.34.

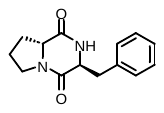
**6m,** cyclo-(D-Pro-L-Phe): fluffy white solid, quant. ^1^H NMR (CDCl_3_, 400 MHz): δ 7.38 – 7.26 (m, 3H), 7.25 – 7.17 (m, 2H), 6.06 (s, 1H), 4.25 – 4.17 (m, 1H), 3.63 (dt, J = 12.4, 8.5 Hz, 1H), 3.40 (ddd, J = 12.2, 9.2, 3.0 Hz, 1H), 3.18 – 3.05 (m, 2H), 3.01 (dd, J = 10.5, 6.5 Hz, 1H), 2.20 (dtd, J = 12.4, 6.6, 1.8 Hz, 1H), 1.94 (dddd, J = 15.1, 7.8, 3.0, 1.8 Hz, 1H), 1.88 – 1.75 (m, 1H), 1.75 – 1.65 (m, 1H).

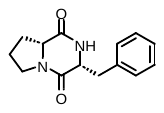
**6n**, cyclo-(D-Pro-D-Phe): yellow-white solid, 83%. ^1^H NMR (CDCl_3_, 400 MHz): δ 7.42 – 7.29 (m, 3H), 7.25 (dd, J = 6.8, 1.9 Hz, 2H), 5.62 (s, 1H), 4.30 (ddd, J = 10.7, 3.9, 1.6 Hz, 1H), 4.10 (ddd, J = 8.5, 6.7, 1.7 Hz, 1H), 3.74 – 3.54 (m, 3H), 2.80 (dd, J = 14.5, 10.7 Hz, 1H), 2.44 – 2.31 (m, 1H), 2.11 – 1.98 (m, 2H), 1.93 (dddd, J = 15.5, 8.8, 7.4, 1.7 Hz, 1H).

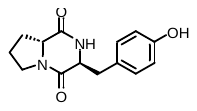
**6o,** cyclo-(D-Pro-L-Tyr): yellow-white solid, 60%. ^1^H NMR (CD_3_OD, 400 MHz): δ 7.07 – 6.94 (m, 2H), 6.76 – 6.67 (m, 2H), 4.14 (td, J = 4.5, 1.0 Hz, 1H), 3.54 (dt, J = 12.0, 8.3 Hz, 1H), 3.37 – 3.25 (m, 6H) *(overlap of 1H with CD_3_OD residual peaks)*, 3.11 (dd, J = 13.9, 4.4 Hz, 1H), 2.88 (dd, J = 13.9, 4.6 Hz, 1H), 2.61 (dd, J = 10.5, 6.2 Hz, 1H), 2.11 – 1.99 (m, 1H), 1.90 (dddd, J = 10.8, 5.5, 4.2, 2.4 Hz, 1H), 1.74 – 1.55 (m, 2H).

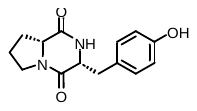
**6p,** cyclo-(D-Pro-D-Tyr): brown-tinted white solid, 26%. ^1^H NMR (CD_3_OD, 400 MHz): δ 7.10 – 7.02 (m, 2H), 6.77 – 6.68 (m, 2H), 4.38 (td, *J* = 4.8, 1.8 Hz, 1H), 4.07 (ddd, *J* = 11.0, 6.4, 2.0 Hz, 1H), 3.57 (dt, *J* = 11.9, 8.3 Hz, 1H), 3.43 – 3.34 (m, 1H), 3.15 – 3.00 (m, 2H), 2.12 (ddt, *J* = 12.1, 6.1, 4.3 Hz, 1H), 1.83 (dddd, *J* = 13.8, 9.8, 5.5, 1.9 Hz, 2H), 1.33 – 1.18 (m, 1H).
